# Species-resolved sequencing of low-biomass microbiomes by 2bRAD-M

**DOI:** 10.1101/2020.12.01.405647

**Authors:** Zheng Sun, Shi Huang, Pengfei Zhu, Lam Tzehau, Helen Zhao, Jia Lv, Rongchao Zhang, Lisha Zhou, Qianya Niu, Xiuping Wang, Meng Zhang, Gongchao Jing, Zhenmin Bao, Jiquan Liu, Shi Wang, Jian Xu

## Abstract

Microbiome samples with low microbial biomass or severe DNA degradation remain challenging for amplicon-based (e.g., 16S/18S-rRNA) or whole-metagenome sequencing (WMS) approaches. Here, we introduce 2bRAD-M, a highly reduced and cost-effective metagenome-sequencing strategy which only sequences ~1% of metagenome and can simultaneously produce species-level bacterial, archaeal, and fungal profiles for low-biomass and highly degraded samples. For mock communities, 2bRAD-M can accurately generate species-level taxonomic profiles for otherwise hard-to-sequence samples with (*i*) low biomass of merely 1 pg of total DNA, (*ii*) high host DNA contamination (99%), and (*iii*) severely fragmented DNA (50-bp) from degraded samples. Tests of 2bRAD-M on stool, skin and environment-surface samples deliver successful reconstruction of comprehensive, high-resolution microbial profiles with agreement across 16S-rRNA, WMS and existing literature. In addition, it enables microbial profiling in formalin-fixed paraffin-embedded (FFPE) cervical tissue samples which were recalcitrant to conventional approaches due to the low amount and heavy degradation of microbial DNA, and discriminated healthy tissue, pre-invasive cancer and invasive cancer via species-level microbial profiles with 91.1% accuracy. Therefore, 2bRAD-M greatly expands the reach of microbiome sequencing.

## Introduction

Metagenome sequencing, widely used to derive the taxonomic profile of microbiome, typically adopts two strategies that target (*i*) amplicons of phylogenetic “marker genes” (e.g., 16S rRNA for bacteria and archaea, and 18S rRNA or internal transcribed spacer (ITS) for fungi) or (*ii*) the whole genomes (whole metagenome shotgun; WMS). Although less costly, marker gene analyses can be limited in taxonomic resolution (i.e. at the genus level) and susceptible to PCR bias in composition and abundance estimates^1^; moreover they are usually unable to capture a landscape-like view that includes bacteria, archaea, fungi and virus due to the lack of universal primers. In contrast, by sequencing the total DNA, WMS can resolve species-or strain-level taxonomy, and offers a landscape-like view that includes all domains of organisms ^2, 3^. However, WMS requires a rather high amount of DNA as the starting material (≥50 ng preferred, 20 ng at a minimum), and are usually unable to tackle DNA samples that are low in biomass, heavily degraded or dominated by host DNA ^1, 4^. Moreover, WMS is typically much more costly, due to the much higher sequencing volume required for covering the whole genome (instead of just the marker gene). Therefore, new methods should be developed that cost-efficiently produce accurate, species-resolution, landscape-like taxonomic profiles for challenging samples like low-biomass, high-host-contaminated and degraded microbiomes.

Restriction site-Associated DNA sequencing (RADseq) that utilizes restriction enzymes to digest genomic DNA from a board range of organisms and sequences only the digested fragments has been applied to genotype variable genetic markers as well as for microbial species detection^5, 6^. Although RADseq has demonstrated the ability to produce species-resolution microbiome profiles with lower costs ^7-10^, the large size variation of DNA fragments after enzyme digestion results in a high bias in PCR-based amplification and thus the low fidelity of reconstructed taxonomic profile^11,12^. Moreover, RADseq has not been thoroughly and systematically benchmarked against marker gene-based or WMS approaches ^7-10^. Therefore, it remains unclear whether RADseq can provide accurate, landscape-like taxonomic profiles for low-biomass and severely degraded microbiome samples.

To address these challenges, here we proposed “2bRAD sequencing for Microbiome” (2bRAD-M) method that utilizes Type IIB restriction enzymes to produce exclusively iso-length DNA fragments of 25-33 bp (depending on the selection of enzymes) for sequencing. This approach reduces the bias in PCR-based fragment amplification, and thus ensures the high fidelity of taxonomic profile. This offers significant benefits especially for low-biomass, heavily contaminated or degraded microbial DNA which requires more PCR cycles. Tests on simulated datasets, mock and actual microbiome samples illustrate that 2bRAD-M, by sequencing just about 1% of genomes, accurately generates species-level taxonomic profiles for challenging samples: (*i*) of merely 1 pg total DNA, (*ii*) of 99% host DNA contamination or (*iii*) consisting of highly degraded fragments just 50-bp long. For real stool, skin and environment-surface samples, it accurately reconstructs a comprehensive, species-resolution profile of bacteria, archaea and fungi. Furthermore, microbiome in the formalin-fixed paraffin-embedded (FFPE) tissue samples which were otherwise recalcitrant to sequencing can now be analyzed, and a species-resolved classifier discriminated the healthy tissue, pre-invasive cancer and invasive cancer with 91.1% accuracy. The ability to profile low-biomass microbiomes at the species level is pivotal to expanding the boundary of known microbial world.

## Results

### The principle and workflow of 2bRAD-M

The principle and appealing features of 2bRAD-M are (**Fig. 1**): (*i*) reliable enzyme-digested sequence tags can be derived that are specific to high-resolution taxa (e.g., species or strain) yet universally applicable for a broad range or all of bacterial, archaeal and fungal genomes; (*ii*) these taxa-specific, iso-length sequence tags can be evenly amplified and sequenced, and (*iii*) the tag sequences can be mapped to reference genomes to reconstruct faithfully the taxonomic composition. Specifically, the experimental workflow has two steps: (*i*) BcgI (a commercially available Type IIB restriction enzymes) is used, as an example, to digest total genomic DNA extracted from microbiome samples. BcgI recognizes the sequence of CGA-N_6_-TGC in the genomic DNA and cleaves on both upstream (12-10 bp) and downstream (10-12 bp) of this signature ^13^, producing short and iso-length DNA (32bp without sticky ends) across all loci ^14,15^. (*ii*) These so-called “2bRAD fragments” are ligated to adaptors, amplified and then sequenced.

**Figure 1.**
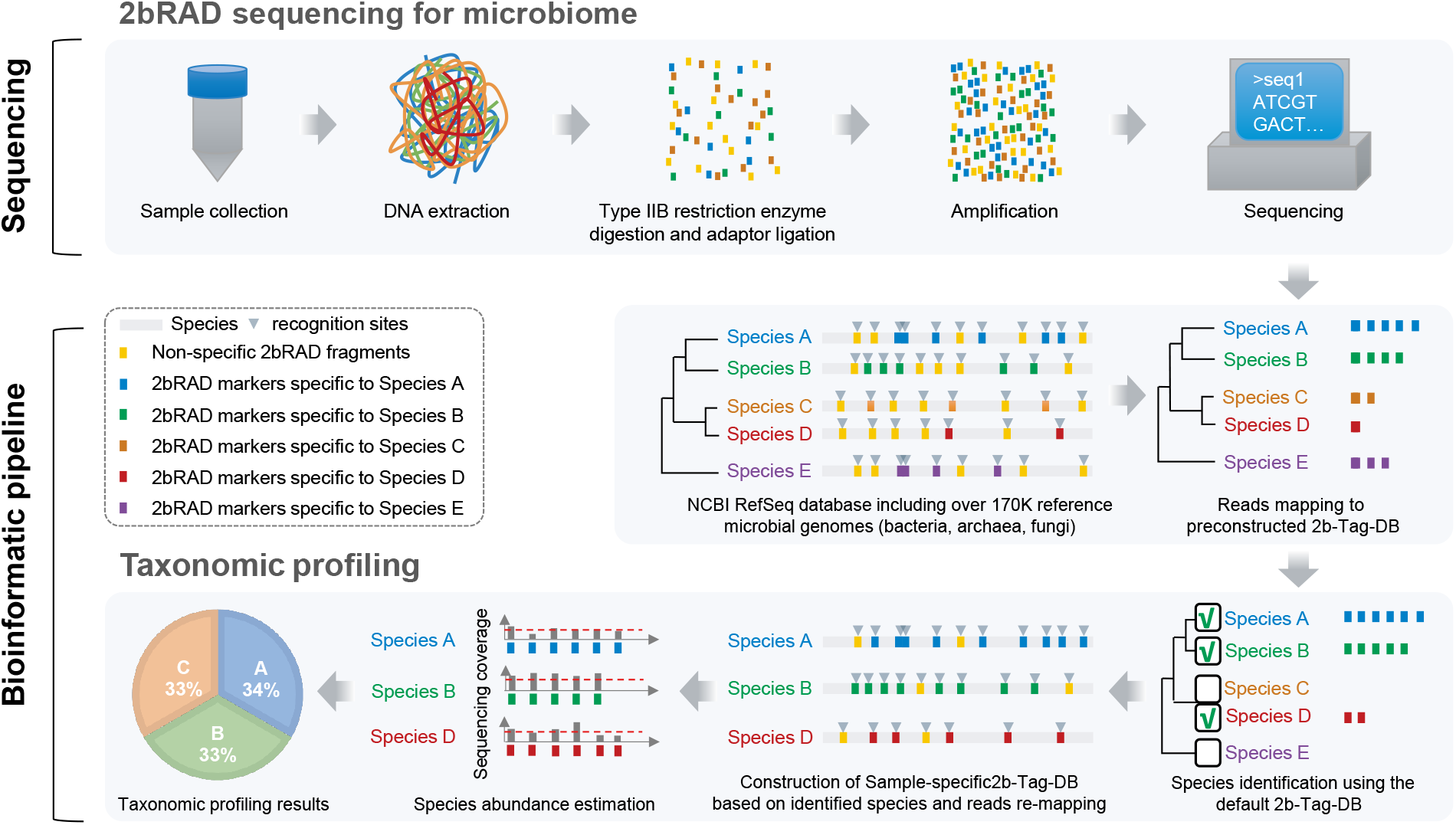
Scheme of the 2bRAD-M workflow. In the library preparation module of 2bRAD-M pipeline, DNA samples were first digested using a type IIB restriction enzyme. The resulting 2bRAD fragments were enriched and amplified for DNA sequencing. In the computational module of 2bRAD-M pipeline, we employed both prebuilt and sample-specific 2bRAD marker database to perform taxonomic profiling on 2bRAD data. Firstly, all reads were mapped against the default prebuilt unique 2bRAD marker database (2b-Tag-DB) to identify all candidate species in a 2bRAD-M sample. Next, to accurately estimate the abundance of identified species, we increased the number of taxa-specific 2bRAD markers for each candidate species by reconstructing a reduced 2bRAD marker database (sample-specific 2b-Tag-DB) which contains more 2bRAD markers specific to each candidate species than those in the default 2b-Tag-DB. All the 2bRAD sequences were then remapped to this sample-specific 2b-Tag-DB for abundance estimation of candidate species. In principle, the relative abundance of a given species was calculated as the read coverage of all species-specific 2bRAD markers. For more information, please refer to **Methods**.

In the computational workflow, the foundation is a unique 2bRAD tag database (“2b-Tag-DB”), which contains taxa-specific 2bRAD tags identified from all the sequenced bacteria, fungi and archaea genomes. Mapping the 2bRAD reads against 2b-Tag-DB thus identifies the presence of species in a sample. Subsequently, to estimate relative abundance of the identified taxa, the mean read coverage of all 2bRAD tags specific to each taxon is derived. To improve utilization rate of reads and classification accuracy, a secondary, sample-specific 2b-Tag-DB was dynamically derived from only those candidate taxa identified in a particular sample, which produces more species-specific 2bRAD tags than the original 2b-Tag-DB and results in more accurate modeling of relative abundance of taxa.

### The feasibility of 2bRAD-M for microbiome analysis via *in silico* simulation

To identify the taxa-specific tags, we first downloaded 173,165 microbial genomes (171,927 bacteria, 293 fungi and 945 archaea; representing 26,163 species) from NCBI RefSeq (Oct, 2019) to create a 2b-Tag-DB via *in-silico* restriction digestion of these genomes using BcgI restriction enzyme. This yields an average of 3,010 iso-length (32bp) 2bRAD tags per genome, and a total of 114,132,487 BcgI-digested unique species-specific 2bRAD tags that are of single-copy within a genome (average 1,194 per species genome; **Supplementary Methods**). Besides BcgI, species-specific 2bRAD tags were also identified for each of the other 15 Type IIB enzymes (AlfI, AloI, BaeI, BplI, BsaXI, BslFI, Bsp24I, CjeI, CjePI, CspCI, FalI, HaeIV, Hin4I, PpiI and PsrI; **Table S1**) to establish the usability of all Type IIB enzymes for 2bRAD-M. Overall, the number of 2bRAD-M tags within a genome and their GC content are highly consistent with the length and the GC% of genomes (*r*>0.98) (**Fig. S1**) ^15^ for all the 16 Type IIB enzymes, suggesting unbiased, broadly applicable representation of the microbial genomes by these tags.

To test whether these species-specific tags enable detection and abundance profiling of all known species in a community, a simulated 50-species microbiome was generated (one genome per species; randomly selected from RefSeq; **Table S2**) and profiled using the default 2b-Tag-DB and sample-specific 2b-Tag-DB for each of the 16 Type IIB restriction enzymes. Performance of 2bRAD-M in species detection was assessed via precision and recall (with relative abundance threshold of 0.0001), while that of species abundance evaluated via L2 similarity score (a metric of similarity adapted from L2 distance ^16^), by comparing to the ground truth (**Fig. 2a**). Precision, recall and L2 similarity of the taxonomic profiling are all remarkably high (average for the 16 enzymes - precision=98.0%, recall=98.0%, L2 similarity=96.9%), and this is achieved with an average genome coverage of 1.50% among the 50 selected genomes (**Fig. 2a**).

**Figure 2.**
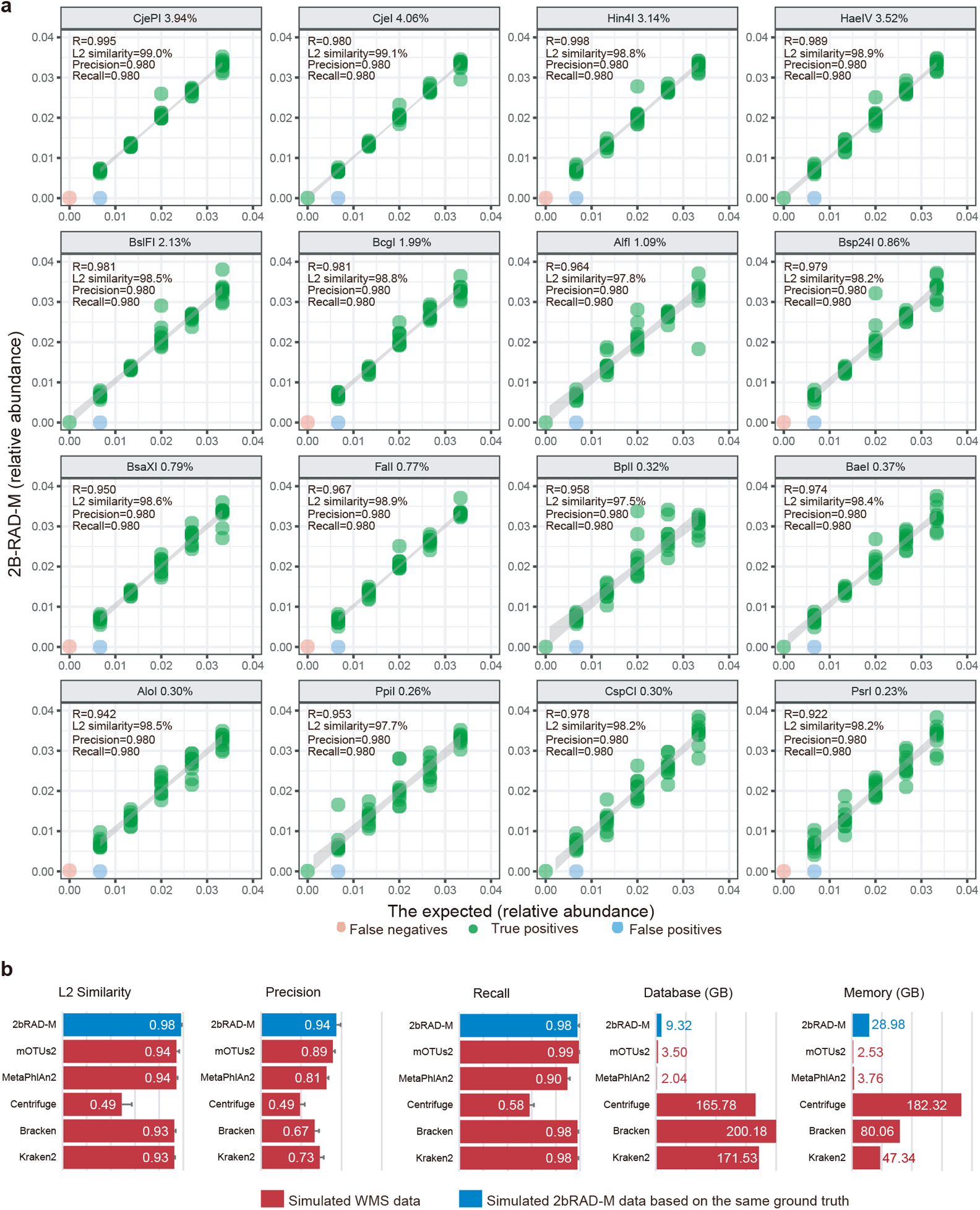
Benchmark measurements of 2bRAD-derived taxonomic profile. (**a**) Simulated microbial community data consisting of 50 microbes were profiled by each of the 16 Type IIB restriction enzymes. The scatter plots indicate the correlation of the taxonomic abundance estimated from 2bRAD-M with the expected abundance for each enzyme. The percentage number indicated in each plot represents the average genome coverage (compare to the original 50 microbial genomes) after digesting by the enzymes. (**b**) Performance comparison of 2bRAD-M with Kraken2, Bracken, Centrifuge, MetaPhlAn2 and mOTUs2 based on five simulated communities.

The accuracy and computational efficiency of 2bRAD-M was assessed by comparison with highly-cited WMS profiling tools such as Kraken^17^, Bracken^18^, mOTUs2^19^ and MetaPhlAn^20^, using the *in silico* simulated data of five gut microbial communities for benchmarking (2bRAD-M applied BcgI-derived species-unique markers to profile the simulated data, while the others produced the abundance profiles from simulated pair-end 150-bp WMS reads). In terms of average precision, recall and L2 similarity (accuracy metrics evaluated based on the ground truth), 2bRAD-M showed a level of 0.94, 0.98, 0.98 respectively, which either outperformed or are equivalent to others (**Fig. 2b**). As for database storage and memory use, 2bRAD-M requires < 10 GB disk space to store the reference marker database, and a relatively low RAM of 30 GB (equivalent to a desktop computer) as compared to Centrifuge, Kraken2 and Bracken (**Fig. 2b**). Thus the 2bRAD-M bioinformatic pipeline can provide accurate profiling results with high computational efficiency.

### High reproducibility and sensitivity of 2bRAD-M under challenging conditions

To assess 2bRAD-M ability to handle low biomass, highly degraded or heavily contaminated samples, we first constructed a mock community (Mock-CAS) consisting of five prevalent oral or gut bacterial species in equal proportion. With this mock community stock, three series of samples were prepared: (*i*) “LoA” (low amount) – samples with total DNA amounts from 50 ng, 20 ng, 10 ng, 1 ng, 100 pg, 10 pg to only 1 pg; (*ii*) “HiD” (high degradation):– 100ng degraded DNA samples where the DNA mixture was randomly sheared by DNAse I into fragments of mostly 150-bp and 50-bp in length; (*iii*) “HoC” (host contamination): – 100ng samples spiked with human DNA to simulate 90% or 99% contamination by host genome. Three technical replicates were included for each of the LoA (n=7∗3), HiD (n=2∗3) and HoC (n=2∗3) groups. Each of the 33 samples from the three series was then sequenced by 2bRAD-M. In addition, a sample of 100ng DNA was then prepared from the mock community stock and sequenced by WMS to be referenced as the positive control (WMS of 50ng or lower DNA failed).

To compare the results and minimize performance bias introduced by bioinformatic pipelines, we employed Centrifuge^21^ to map both WMS (150-bp) and 2bRAD-M (32-bp) generated reads to the reference microbial genomes for taxonomic abundance estimation. For 2bRAD-M, tags from the actual data of each sample covered avg. 97.1% *in silico* predicted tags from the five genomes, indicating high consistency between observed and expected 2bRAD tags (which is key to accurate and reliable taxonomic profiling). Moreover, for each sample, the 2bRAD-M results are highly consistent among the three biological replicates (avg. L2 similarity: 95.4%; **Fig. 3a**).

**Figure 3.**
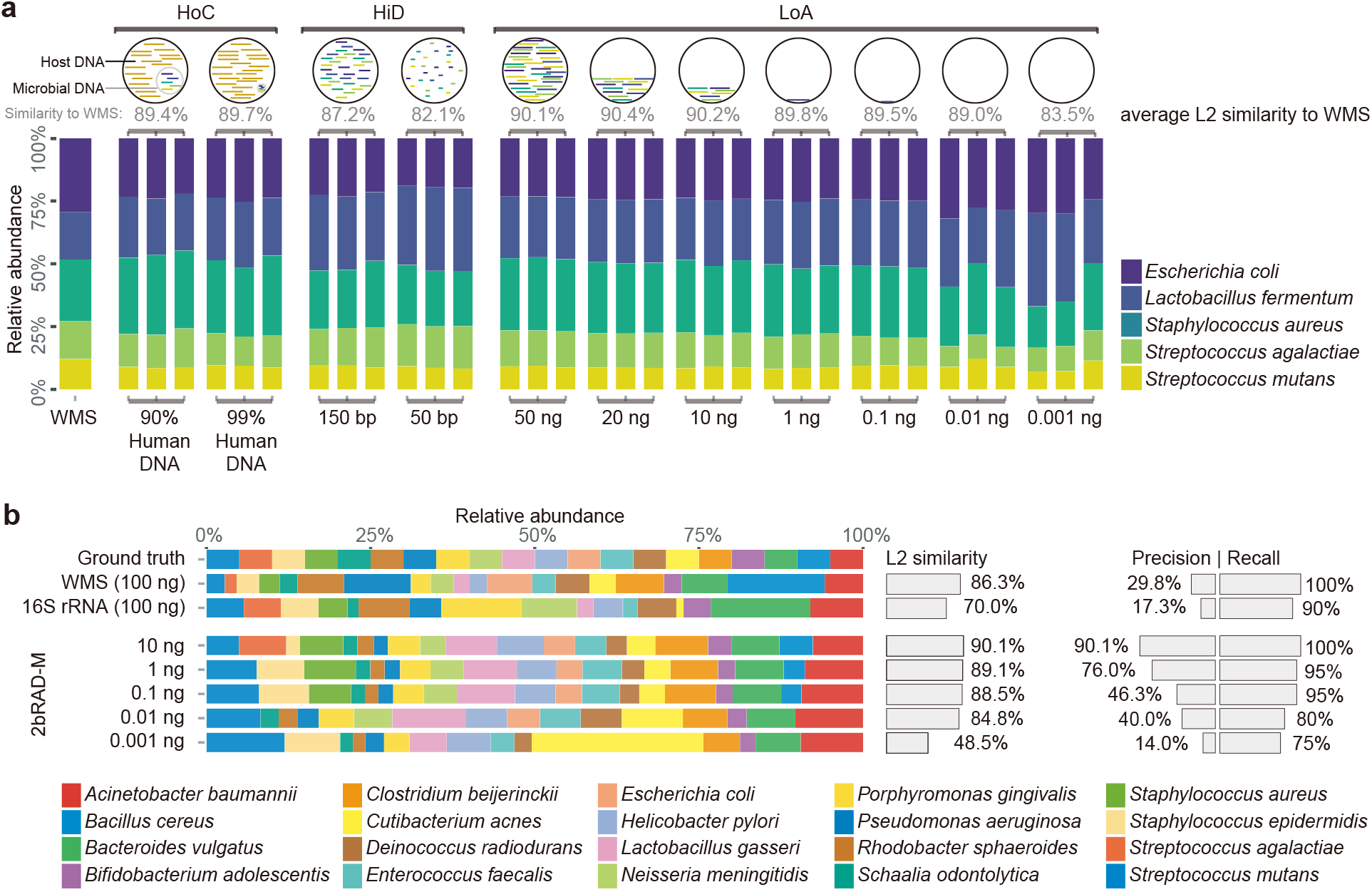
Comparison of taxonomic profiling results of 2bRAD-M, shotgun WGS and 16S rRNA sequencing methods in mock microbial communities. (**a**) 2bRAD-M performance in samples with low amount (LoA), high DNA degradation (HiD) and high host DNA contamination (HoC) based on mock community of five bacteria species (Mock-CAS). LoA are samples with a gradient of total DNA concentrations (50 ng, 20 ng, 10 ng, 1 ng, 0.1 ng, 0.01 ng, and 0.001 ng); HiD includes samples with fragmented bacterial DNA (50bp or 150bp). HoC are samples with mixture of human DNA (90% or 99%) and bacterial DNA. Three technical replicates were included for each group. The L2 score that measures the microbial composition similarity between 2bRAD-M and WMS is shown on the head of each stacked bar plot. The false-positive identification rates of reads (i.e., reads mapped to species not in the mock community) are very low (0.9% in WMS and 1% in 2bRAD-M). (**b**) Benchmark of WMS, 16S rRNA and 2bRAD-M approaches based on ATCC MSA1002 mock community. Each stacked bar plot in the left panel shows the resultant taxonomic profile of a given metagenomic method prepared at a specified DNA amount. The right panel indicates the corresponding L2 similarity, precision and recall score as compared to the ground truth.

Notably, for the LoA, HiD and HoC groups, average L2 similarity between 2bRAD-M profiles and the reference positive control is 88.9%, 84.6% and 89.6% respectively, with global average being 88.2% (ranging from 82.1% to 90.4%; **Fig. 3a**). These observations are consistent across the technical replicates, indicating high reproducibility of 2bRAD-M. Specifically, in the LoA group, the L2 similarity of the 1 pg sample can still realize a respectable 83.5% as compared to 90.1% for the 50 ng sample (**Fig. 3a**). This suggests that 2bRAD-M offers high sensitivity and stable performance in low biomass samples over a broad DNA-amount range (from 50 ng to 1 pg). In the HiD group, the L2 similarity is 87.2% and 82.1% for the 150bp- and 50bp-samples respectively, indicating DNA degradation did not have a large negative effect and the 2b-RAD-M can effectively accommodate severe DNA degradation while providing reliable results. The L2 similarity for the HoC group is 89.4% in the 90% host-contaminated samples and 89.7% in 99%-host-contaminated samples, suggesting 2bRAD-M’s ability to provide reliable microbial profiling in the presence of host-DNA contamination.

To further probe 2bRAD-M proficiency in profiling highly complex microbial community at low biomass setting, we used ATCC MSA1002, a standard, commercially available mock community consisting of 20 bacterial species (from 18 genera) with at equal DNA abundance among species^22^. Samples with total DNA amount range from 10 ng, 1 ng, 100 pg, 10 pg and 1 pg were prepared from this mock for 2bRAD-M profiling. As a reference control, a 100 ng sample was profiled with 16S-rRNA and WMS approach (the 1pg ~10ng samples failed). The precision for 2bRAD-M is at 90.1%, 76.0%, 46.3%, 40.0% and 14.0% for the 10 ng, 1 ng, 100 pg, 10 pg and 1pg samples respectively; this is in contrast to the 29.8% of WMS-100ng and the 17.3% of 16S-rRNA-100ng (**Fig 3b**). This demonstrates that 2bRAD-M offers much lower false positives in detecting the species than 16S-rRNA and WMS. For recall, 2bRAD-M exhibits 100.0%, 95.0%, 95.0%, 80.0%, 75.0% for 10 ng, 1 ng, 100 pg, 10 pg and 1 pg respectively, as compared to the 100.0% of WMS-100ng and the 90.0% of 16S-rRNA-100ng. Therefore, 2bRAD-M offers comparable level of sensitivity as WMS and 16S-rRNA with down to 100 pg of DNA amount. The L2 similarity for 2bRAD-M is 90.1%, 89.1%, 88.5%, 84.8% and 48.5% for 10 ng, 1 ng, 0.1 ng, 10 pg and 1pg respectively, as compared to 86.3% for WMS-100ng and 70.0% for 16S-rRNA-100ng. Taken as a whole, these results evidently demonstrated 2bRAD-M’s ability to profile complex microbiota with a high level of sensitively and specificity.

### 2bRAD-M enables cost-effective deep microbiome profiling of real samples (fecal)

To assess the performance of 2bRAD-M on real samples, we performed and compared 2bRAD-M and 16S rRNA sequencing on human fecal samples (n=3). In addition, each sample was also subjected to ultra-deep WMS sequencing (mean of 437 million reads or 239.73 Gb per sample) with the resultant taxonomic profiles used as evaluation reference^23^.

Taxonomic profiles from 2bRAD-M are compared with those from 16S rRNA sequencing (**Fig. 4a-c**) and WMS (**Fig. 4d-f**; **Supplementary Methods**). Specifically, at the genus-level, results of 2bRAD-M and 16S rRNA are highly consistent (mean Pearson correlation R=0.997 and mean L2 similarity L2=92.0%), and avg. 95.27% of genus identified by 16S were also detected by 2bRAD-M (**Table S3**). Importantly, at the species level, 2bRAD-M and the WMS are also concordant, as evidenced by a high Pearson correlation (R=0.974) and high L2 similarity (up to 88.6%). Moreover, only 3.35% of the taxa in WMS were not identified in 2bRAD-M. Thus 2bRAD-M can produce highly complete and accurate species-level profiles that are equivalent to WMS.

**Figure 4.**
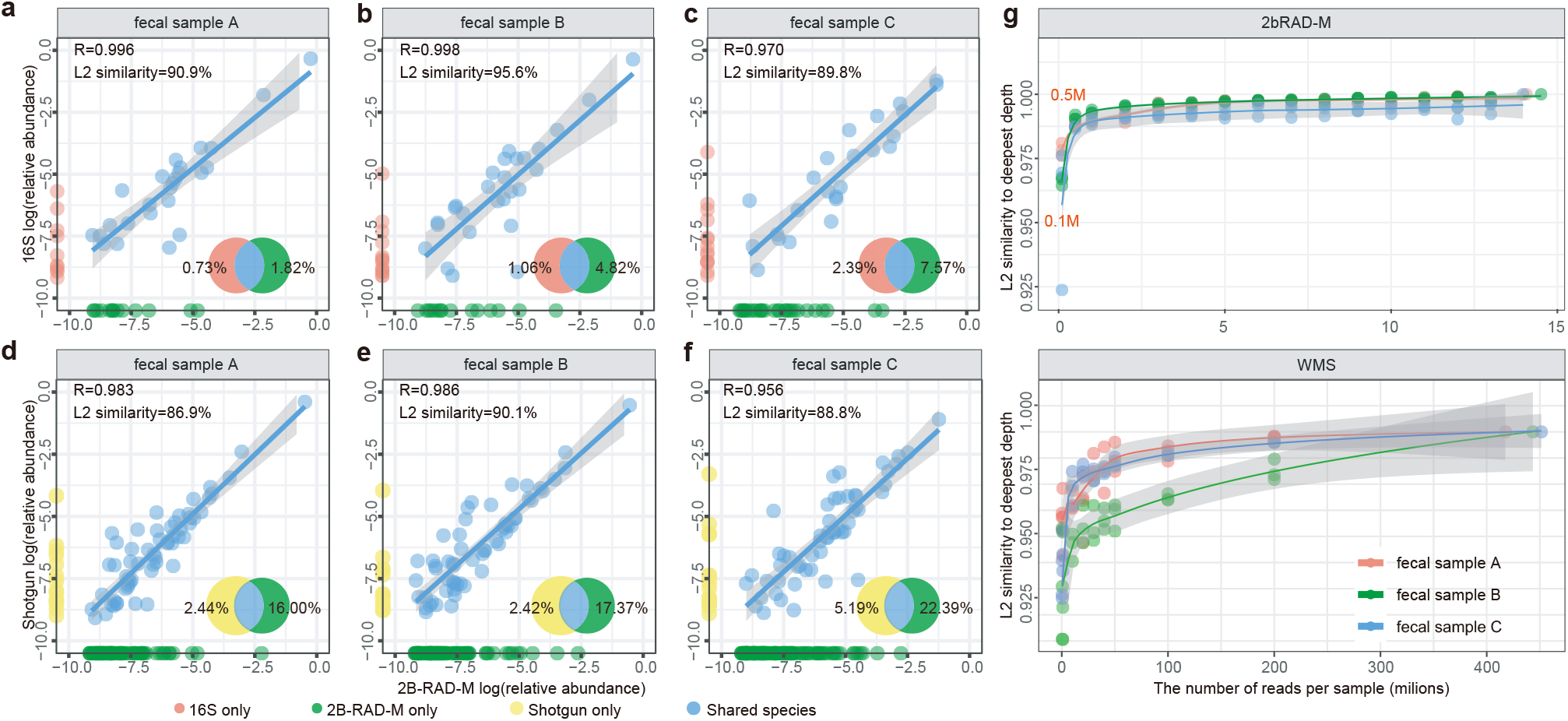
Comparison of taxonomic profiles and desired sequencing depth of 2bRAD-M, 16S and WMS in fecal samples. (**a-c**) Comparison of taxonomic profiles at the genus level between 16S rRNA and 2bRAD-M. (**d-f**) Comparison of taxonomic profiles at the species level between WMS and 2bRAD-M. (**g**) The rarefaction analysis of 2bRAD-M and WMS samples. The species-level compositions in the subsampled data at each given sequencing depth were compared to the pre-rarefaction result for each method.

One determinant of quality (and cost) of taxonomic profiling is sequencing depth ^24^. To probe how it influences 2bRAD-M performance, the fecal sequencing datasets were subsampled to various depths for 2bRAD-M (average 14 million reads per sample, subsampled at 0.1, 1, 2, 3, 5 to 14 million sequences per sample) or WMS (average 437 million reads per sample, subsampled at 0.5, 10, 20, 30, 40, 50, 100, and 200 million sequences per sample) respectively. Then we asked how many 2bRAD sequences are required to accurately quantify key ecological metrics such as alpha diversity (Shannon index), beta diversity (Bray-Curtis), and species compositions (species richness). In each of the fecal samples, with 20-40 million reads per sample, WMS identified averagely 170-182 species, but the Shannon diversity and species abundance estimates are far from saturation (i.e., reaching the values derived by sequencing ~400 million reads). In contrast, with just 3-4 million reads per sample, 2bRAD-M identified 173-188 species in each fecal samples and yield Shannon diversity and species abundance estimates comparable with those from the deepest (~14 million) read depth (**Fig. S2**).

To evaluate the optimal sequencing depth of 2bRAD-M for species-level taxonomic profiling, we evaluated the similarity of taxonomic profiles derived at each reduced sequencing depth to its original taxonomic profile using L2 similarity score. For 2bRAD-M, a sequencing depth of over 0.5 million reads per samples can achieve a L2 similarity score of 98.9% (1 million for 99.1%), which however requires many more (at least about 200 million) reads per sample in WMS (**Fig. 4g**). Overall, with ~3 million reads (i.e., 60 Mb of sequencing data) per sample, 2bRAD-M can generate consistent, accurate and stable alpha diversity estimates and taxonomic profile at the species level.

### 2bRAD-M enables species-resolved analysis of low-biomass skin, home and car samples

To assess 2bRAD-M performance on actual low-biomass samples, we collected samples from human skin surface (underarm, n=20; **Table S4**) for 16S-rRNA and 2bRAD-M analysis (WMS was aborted due to too-low DNA amounts for library construction). To gauge the potential impact of contaminating DNA (introduced from regents, in workflow, etc.) in the low-biomass samples ^25^, MSA1002 was included as a control. MSA1002 profiling revealed the expected microbial community profile while unforeseen microbes were absent which indicates minimal contamination. Between 16S rRNA and 2bRAD-M, a high degree of consistency in taxonomic profiles at the genus level was observed, with avg. L2 similarity of 81.1% (**Table 1**; **Fig. S3**). In both methods, *Staphylococcus* (2bRAD-M: avg. 47.17%; 16S rRNA: avg. 44.43%) and *Corynebacterium* (2bRAD-M: avg. 10.38%; 16S rRNA: avg. 15.06%) are recognized as dominant microbes, consistent with existing literature ^26,27^. Interestingly, unsupervised clustering at genus-level uncovered two distinct clusters among the 20 underarm samples, with one featuring higher abundance of *Staphylococcus* observed in both 16S-rRNA and 2bRAD-M (2.41-fold, *p*=4.6E-06, Wilcoxon rank-sum test, for 16S rRNA; 3.20 fold, *p*=1.08E-05, Wilcoxon rank-sum test, for 2bRAD-M; **Fig. 5a**). Notably, the species-level profile from 2bRAD-M readily revealed that the observed difference in *Staphylococcus* was defined by higher relative abundance of *S. epidermidis* (4.63-fold, *p*=4.3E-04), with no significant change in other *Staphylococcus* spp. between the clusters (**Fig. 5b**), which suggests interesting biological hypothesis on *S. epidermidis* functionality (notably, *S. epidermidis* can protect against pathogens such as *S. aureus* by producing antimicrobial peptides that modulate skin microbiome dysbiosis in atopic dermatitis ^28^). These observations would have been missed with 16S rRNA, thus 2bRAD’s ability to resolve such species-level heterogeneity (e.g., *Staphylococcus spp*.) in low-biomass samples can potentially enable novel biological insights.

**Table 1.** High concordance of dominant microbial taxa in 16S rRNA and 2bRAD-M profiling results of 20 underarm samples.

| Top genus in 16S | Relative abundance | Rank | Corresponding species in 2bRAD-M | Relative abundance | Rank |
| --- | --- | --- | --- | --- | --- |
| *Staphylococcus* | 44.43% | 1 | *Staphylococcus epidermidis* | 27.56% | 1 |
|  |  |  | *Staphylococcus hominis* | 12.57% | 2 |
|  |  |  | *Staphylococcus capitis* | 4.78% | 5 |
|  |  |  | *Staphylococcus haemolyticus* | 1.78% | 11 |
|  |  |  | The other 4 species | 0.48% |  |
|  |  |  | SUM | 47.17% |  |
| *Corynebacterium* | 15.06% | 2 | *Corynebacterium ureicelerivorans* | 2.58% | 8 |
|  |  |  | *Corynebacterium jeddahense* | 2.41% | 9 |
|  |  |  | *Corynebacterium aurimucosum* | 1.80% | 10 |
|  |  |  | *Corynebacterium pseudogenitalium* | 1.29% | 15 |
|  |  |  | *Corynebacterium kefirresidentii* | 1.23% | 17 |
|  |  |  | *Corynebacterium tuberculostearicum* | 1.08% | 19 |
|  |  |  | The other 47 species | 2.08% |  |
|  |  |  | SUM | 12.47% |  |
| *Uruburuella* | 5.67% | 3 | *Uruburuella suis* | 5.80% | 4 |
| *Enterococcus* | 4.49% | 4 | *Enterococcus devriesei* | 1.68% | 12 |
|  |  |  | *Enterococcus dispar* | 1.48% | 13 |
|  |  |  | The other 3 species | 0.45% |  |
|  |  |  | SUM | 3.61% |  |
| *Moraxella* | 02.34% | 5 | *Moraxella osloensis* | 2.77% | 7 |
|  |  |  | The other 1 species | 0.04% |  |
|  |  |  | SUM | 2.81% |  |
| **SUM** | **72.00%** |  | **SUM** | **71.86%** |  |

**Figure 5.**
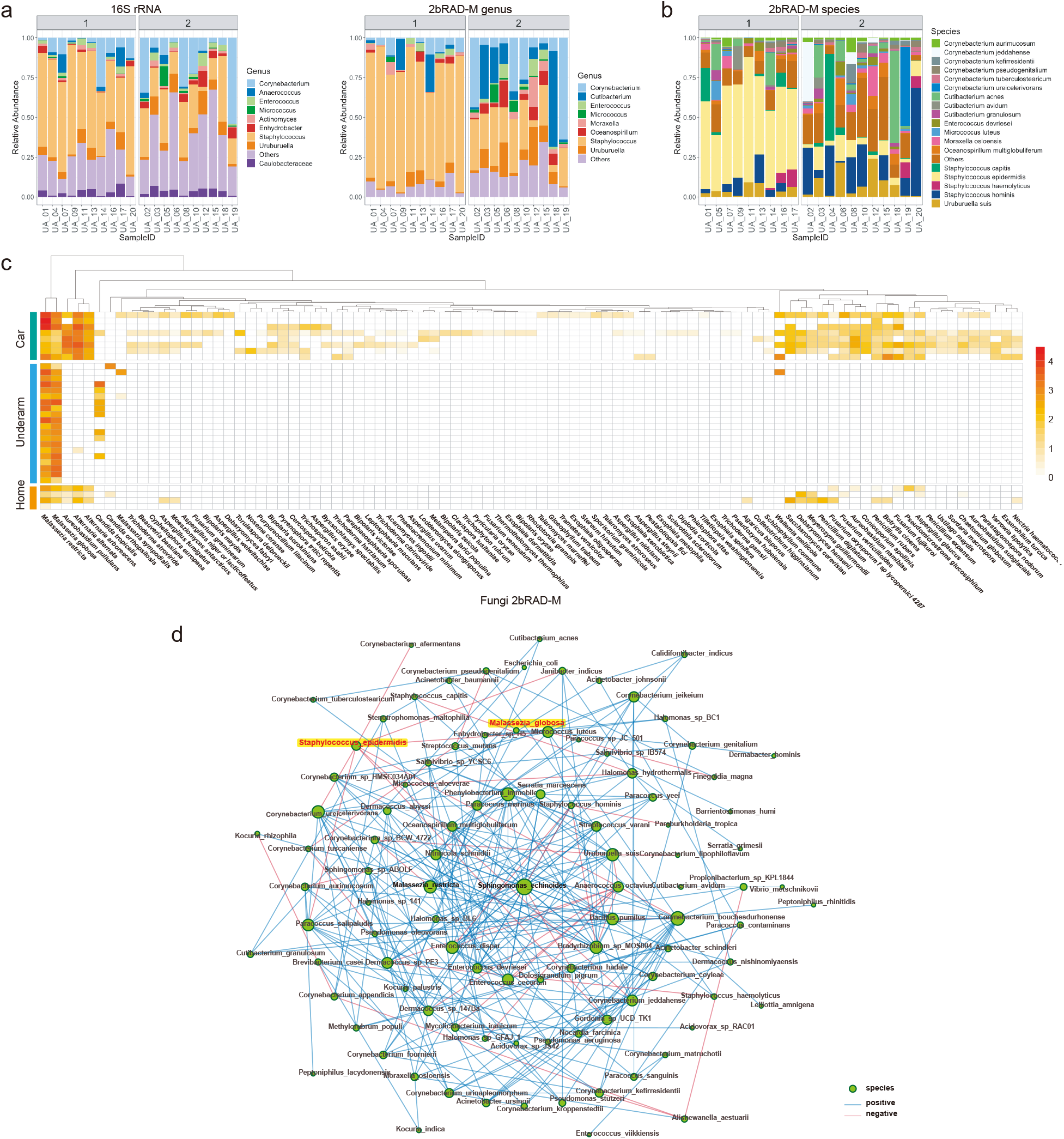
2bRAD-M analysis of low biomass samples collected from underarm, home and car surfaces. (a) Relative abundance of bacteria at genus-level (>1%) identified in underarm by 16S rRNA sequencing (left panel) and 2bRAD-M (right panel) with the samples stratified by cluster group. (**b**) Relative abundance of bacteria at species-level taxa (>1%) identified in the underarm by 2bRAD-M with the samples stratified by cluster group. (**c**) Heat map shows identified fungal species via 2bRAD-M in the underarm, home and car samples. (**d**) Co-occurrence network of bacterial and fungal species based on 20 underarm samples profiled by 2bRAD-M. Each green circle represents a species, and its size refers to the degree centrality score. Blue/red edges indicate positive/negative spearman coefficients.

We further explored the application of 2bRAD-M, by testing on in-door environmental low-biomass samples, which are of significant health implications due to their contact with humans^29^. Surfaces of floor mat (n=5) and cushion (n=3) in car, and interior surfaces of a house (n=4) were sampled and subjected to both 16S rRNA and 2bRAD-M (similarly, WMS was aborted due to the low DNA amounts; **Table S4**). High L2 similarity (average 85.6% among samples; **Fig. S3**) between 2bRAD-M and 16S rRNA was observed, suggesting stable performance of 2bRAD-M for such in-door samples. At the species level (**Table S5**), we identified 4929, 2335 and 1626 taxa for the three niches respectively, whose dominant bacterial species are highly distinct: (*i*) floor mats are dominated by *Kocuria rosea* (23.23%), *Psychrobacter 1501_2011* (11.44%) and *Acinetobacter johnsonii* (10.43%); (*ii*) cushions are dominanted by *Cutibacterium acnes* (42.19%), *Lactobacillus delbrueckii* (10.62%) and *Ralstonia pickettii* (10.14%); (*iii*) home surfaces are mainly colonized by *Streptococcus mitis* (49.64%), *Prevotella copri* (39.14%) and *Megamonas funiformis* (18.36%). Rarefaction of sequence depth (e.g., sequenced 2b-tags) via alpha diversity, beta diversity and species-level compositions supports high robustness of 2bRAD-M for such low-biomass in-door samples (**Fig. S4**).

Similar to WMS, 2bRAD-M enabled species-level profiles for fungi along with bacteria. In general, the relative abundance of fungi in the underarm and indoor environment samples is extremely low (0.83%) as compared to bacteria (99.16%) (**Table S6**). Possible reasons for this observation include: (*i*) micro-environments of these samples are not friendly fungal growth^30^; (*ii*) the relatively small number of available fungi genomes curated in our reference database limits the discovery of the fungi population. Nonetheless, among all the sites, the samples taken from car cushion are found to harbor the highest amount of fungi (2.33%) with the surfaces from home being the lowest (0.06%). In addition, the 2bRAD-M species-level profile unveils distinctive patterns of fungal composition among the various sites (**Fig 5c**). *Malassezia restricta* and *Malassezia globose*, known commensals on human skin surface^31^, were found as the top abundance fungi at underarm and most of the indoor environmental samples. It was possible that the *Malassesia* sp. was transferred from human, given the likely high contact frequency between the skin and these indoor environmental surfaces. Conversely, *Alternaria alteranta* was mostly found in the indoor environments but absence in the underarm which is in agreement with existing reports^32, 33^. Thus the fungi profiles derived from 2bRAD-M are consistent with literature, demonstrating the ability to reliably profile fungi (simultaneously with bacteria) in low-biomass samples.

Taking advantage of 2bRAD-M’s capability in profiling bacterial and fungal species simultaneously, occurrence-network analysis revealed negative correlation between the human skin commensal yeast of *Malassezia globosa* (which is associated with Seborrheic Dermatitis ^34^) and the aforementioned *S. epidermidis* (recently proposed as a gatekeeper of healthy skin ^35^; Spearman coefficient of -0.569; **Fig. 5d**). Such landscape-like, species-level correlations for low-biomass microbiomes can potentially reveal novel bacteria-fungi interactions.

### 2bRAD-M enables tumor microbiome profiling from FFPE tissue samples

Microbiota in human tumor or blood tissues were recently associated with the types, developmental stages or chemotherapeutic efficiency of cancer ^36-40^. Formalin-fixed, paraffin-embedded (FFPE) tissue, the gold standard of preserving tumor biopsy specimens ^41, 42^, represents a vast, irreplaceable historical clinical resource of enormous value for cancer microbiome studies^43^, however profiling microbiome from FFPE tissues has been challenging due to the low microbial biomass, high human DNA contamination, severe DNA damage and cross-links by chemical modification ^44^. Therefore 2bRAD-M is an ideal approach to elucidate the microbial community in FFPE samples. Here, we collected FFPE cervical tissue samples from 15 healthy controls (H), 15 pre-invasive cancerous (PreC; benign) and 15 invasive cancerous patients (InvaC; malignant), and subjected these samples to 2bRAD-M sequencing. DNA from the FFPE tissue was extracted from an area of 3cm^2^ with 4μm thickness. On average 25 ng/ul of DNA in each sample was detected as smears of size under 500 bp in agarose gels (**Fig. S5**), suggesting extremely low concentration and highly fragmented nature of DNA (both human and microbes ^44^) in FFPE samples. The microbiome of the FFPE samples is mostly dominated by bacteria species (n=243) with minimal fungal species (n=2; **Table S5**). The alpha diversity (Shannon and Simpson index) in healthy controls (H) is significantly lower than PreC and InvaC (*p*=0.044, Kruskal test; **Fig. 6a**). Among the identified bacterial species (**Fig. S5**), samples in the PreC and InvaC groups are significantly enriched with *Methyloversatilis discipulorum* (*p*-value=1.2e-5), *Mycobacterium tuberculosis* (*p*-value=0.004), *Methyloversatilis universalis* (*p*-value=1.8e-5), *Ferrovibrio K5* (*p*-value=0.001) and *Pseudomonas aeruginosa* (*p*-value=6.8e-5) (each increase in relative abundance from H to PreC and to InvaC; **Fig. 6b**). Conversely, *Lactobacillus* spp. (*L. paracasei, L. vaccinostercus*,, *L. pentosus* and *L. plantarum*) are greatly enriched in H (average abundance of 63.1%, in contrast to 33.9% and 32.9% in PreC and InvaC respectively, **Fig. 6b**); this is consistent with a previous study^40^ showing depletion of the *Lactobacillales* genera in “fresh”, non-FFPE cervical cancerous tissues. Notably, *Lactobacillus paracasei* shows anticancer potential against cervix cancer cells (HeLa) *in vitro*^45^. Furthermore, the enrichment of *Mycobacterium tuberculosis, Pseudomonas aeruginosa* and *Staphylococcus aureus* in PreC- and InvC-phase PPFE samples is consistent with “fresh” tissue based studies that reported related genus-level (via 16S ^32^) or species-level (via WMS ^40, 46^) taxa. Taken as a whole, 2bRAD-M has successfully captured the microbiome structure in FFPE samples, and is able to reveal previously unknown discriminative microbial features between healthy and cancerous tissues. These features may serve as potential indicative novel markers for tumor onset and progression diagnosis. To evaluate this potential, we applied Random Forest on the FFPE taxonomic profiles at the species-level, and the model distinguishes H, PreC and InvaC samples with 91.1% accuracy (ten-fold cross validation; **Fig. 6c**). Notably, we can achieve maximized discriminative performance (AUC: 0.96) with the fewer features of the nine most important species in the RF model (**Fig. 6d**; **Table S7**). Thus, 2bRAD-M offers a viable option for microbiome profiling on the vast archive of historical FFPE sample with potential application in early diagnosis and treatment of cancer.

**Figure 6.**
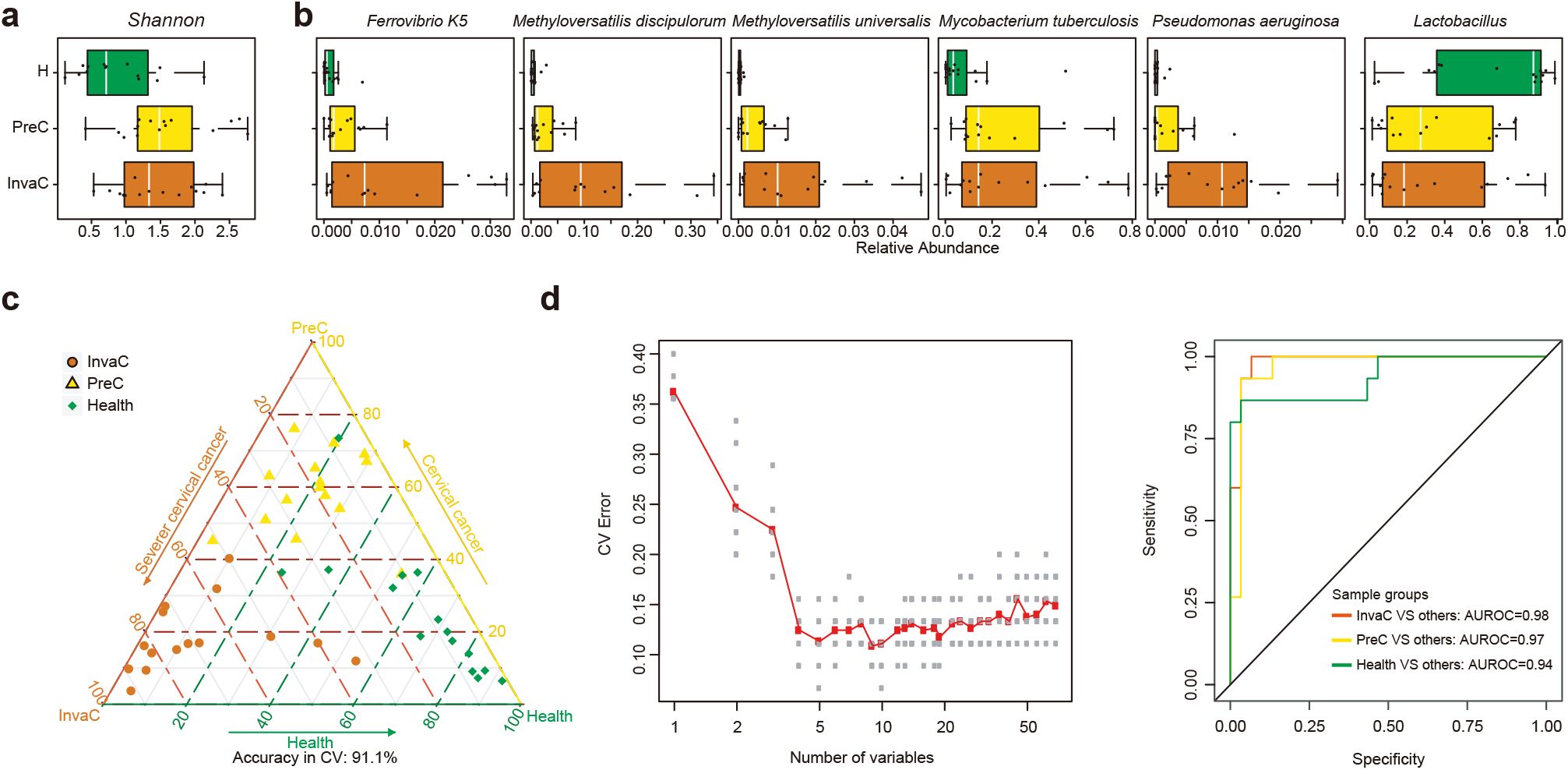
Microbiome-based diagnosis of FFPE thin section from cervical cancer samples as enabled by 2bRAD-M. (**a**) Shannon and Simpson index among 15 healthy controls (H), 15 pre-invasive cancerous (PreC; benign) and 15 invasive cancerous (InvaC; malignant) samples. (**b**) Comparison of differential species and *Lactobacillus spp*. among the three groups. (**c**) The Random Forest classifier for discriminating cancer and healthy samples. In the ternary plot, each dot represents a FFPE sample. The axes indicate the microbiome-based probability of being InvaC, PreC and H for a FFPE sample. The closer one sample is to an apex, the more likely it is predicted as to be corresponding disease states. (**d**) Feature selection by rebuilding Random Forest classifiers using a series of reduced sets of features. The scatter plot shows that nine variables (species) in a reduced RF model (i.e., the AUC plot on the right) can maximize model performance. And the ROC curve shows an even better discriminate performance using binary categories (averagely 0.96).

## Discussion

Using mock microbiomes plus actual samples from stool, skin, environment-surface and frozen FFPE tissues, we demonstrated the ability of 2bRAD-M to profile microbiome samples that are challenging by using marker-gene or WMS sequencing approaches. Three key features are highlighted from this research. *Firstly*, it can analyze samples with low biomass (down to 1 pg), severe degradation or high contamination. This advantage is built on the ability to (*i*) provide an unbiased (i.e., without the large size variation of DNA fragments) and reduced representation of metagenomes for sequencing, (*ii*) evenly cover all restriction sites, and (*iii*) generate short, iso-length reads and thus reliable quantification of tag abundance. With such advantage, 2bRAD-M is valuable to profiling microbiome samples from sparsely populated niches (e.g., indoor surfaces, blood and skin), precious clinical specimen or heavily degraded or contaminated tissues (e.g., FFPE sections and archaeological samples). Notably cervical cancer can be effectively cured in the preinvasive period ^47^, yet detection of this asymptomatic stage remains challenging which results in unnecessary delays in diagnosis and treatment ^48^; therefore, 2bRAD-M sequencing of FFPE cervical cancer microbiomes, which is challenging for WMS or 16S rRNA approaches, promises to turn the enormous clinical repositories of FFPE tissue specimens into a treasure trove of microbiome-driven discoveries ^41^.

*Secondly*, it can provide species-level taxonomic information for bacteria, archaea and fungi simultaneously. The multiple species-specific 2bRAD-M tags can uniquely identify the species from the microbial community, as resolution at the “species” level instead of the “genus” level can be crucial ^49^. In this case, the ability to extract the species-level cancer-stage-specific markers from FFPE tissues, as well as to distinguish *S. epidermidis, S. hominis* and *S. aureus* from underarm skin samples, are crucial to diagnosis and mechanistic understanding of inflammation ^28^. Interestingly, the overall ratio of fungal and bacterial abundance, a potential indicator of ecological balance^50^, is 0.31% for underarm, 0.07% for home and 1.27% for car (significantly different), suggesting characteristic ecological features among the sites.

*Finally*, using 2bRAD-M, the species-level taxonomic profiling can be achieved with a much lower sequencing cost than WMS. For example, only 1~5% of the sequencing data of WMS are required by 2bRAD-M to produce a taxonomic profile of equivalent accuracy, resulting in a cost reduction of 20-100 folds. Moreover, multi-isoRAD allows sequencing of five concatenated 2bRAD tags altogether via a single, 100–150 bp read from Illumina paired-end (PE) library ^14^, which further reduces the sequencing cost. The cost advantage is significant when high sequencing depth is required to address questions like unveiling the true microbial diversity or recovering rare but functionally important species in complex microbiota ^24, 51^. Thus 2bRAD-M appears especially suitable for such samples or circumstances.

Notably, the performance of 2bRAD-M is limited by the availability and the potential bias of related reference genomes (e.g., 171927 bacterial and 945 archaeal genomes, yet only 293 fungal genomes in RefSeq), although the situation is rapidly improving with the steady increase of sequenced genomes for both cultured and uncultured microbes (e.g. via single-cell sequencing ^52^). With the key advantages versus marker genes analysis and WMS, 2bRAD-M is particularly suitable for cost-effectively deriving a species-resolution taxonomic profiling for samples with low biomass (down to 1 pg), severe degradation or high contamination, thus greatly expands the opportunities in microbiome research.

## Materials and Methods

### Sample preparation and sequencing

Two mock microbial communities were used to validate the stability, sensitivity and precision of 2bRAD-M. The first consists of five evenly mixed bacterial strains including *Streptococcus mutans* UA159, *Streptococcus agalactiae* ATCC13813, *Staphylococcus aureus* ATCC29213, *Escherichia coli* DH5α and *Lactobacillus fermentum* ATCC9338. Then three circumstances that simulated “challenging” microbiome samples were produced (with three replicates for each sample): (1) LoA: samples with low amount DNA; A concentration gradient from 10ng to 1pg was designed, with one tenth of the concentration retained each time. (2) HiD: samples with highly degraded DNA; Two samples with DNA length about 150bp and 50bp were included. (3) HoC: samples with host DNA contamination, which consist of either 90% or 99% human DNA.

The second mock used in this paper is the 20-Strain Even Mix Genomic Material (named as Mock MSA1002, 3.87 ng/uL; frozen 50 µL in Tris-HCl pH 8.5) which is purchased from ATCC. This mock comprises genomic DNA prepared from fully sequenced, characterized, and authenticated ATCC Genuine Cultures that were selected based on relevant phenotypic and genotypic attributes, such as Gram stain, GC content, genome size and spore formation.

#### Fecal and FFPE tissue samples

The study protocol complies with the ethical guidelines of the 1975 Declaration of Helsinki and was approved by the DOST ethical committee and conducted according to ICH guidelines for Good Clinical Practice. Written informed consent and photography consent were obtained from each subject before enrollment. Three healthy adult individuals were enrolled as volunteers and fecal samples were collected for deep WMS, 16S rRNA gene amplicon and 2bRAD-M sequencing for comparison.

Totally 45 cervical cancer-related sections in FFPE blocks (with a thickness of 4μm and an area of 3cm^2^) underwent microbiome profiling via 2bRAD-M. The original tissue samples include 15 healthy controls (H), 15 pre-invasive cancerous (PreC; benign) and 15 invasive cancerous (InvaC; malignant) ones (determined by morphological evidence of polymorphonuclear infiltration). Each of the 45 samples was from a distinct individual, i.e., a cross-sectional design. To prepare the FFPE blocks, fresh tissue samples were fixed in phosphate buffered formalin for 24–48h, followed by tissue processing for 10h, and paraffin embedding for 20min ^53^. Post-fixation processing of the tissues was completed in a histopathological laboratory using consistent processor protocols over years. Prior to microbiome profiling, the FFPE blocks were already stored at 17°C –22°C and 20%– 60% humidity levels for 1-2 years. For these FFPE tissue sections, attempts to construct 16S amplicon or WMS libraries for microbiome sequencing both failed, due to the low quality of initial DNA (**Fig. S6**).

#### Underarm, home and car sampling collection protocol & processing

For the underarm sampling protocol, individuals were subjected to a washout period where the use of anti-bacterial products were not allowed for 4-5 days. After the washout period, 22mm D-squame tape strip (Cuderm) was applied onto the lower underarm skin surface (without hair) using a pressure applicator. The tape strips were then pre-treated with 200 mg of 0.1 mm Zirconia Silica beads and bead beat using Qiagen TissueLyser II (Valencia, CA) at 30 Hz for 3 minutes to lyse (via mechanical force) and dislodge the biomass from the tape strip. After dislodging the biomass from tape strip, complete cells lysis is achieved via further enzymatic reaction by 2% w/v lysozyme, 0.05% w/v lysostaphin and 1.2% Triton-X in TE buffer, followed by DNA extraction using Qiagen DNeasy Blood and Tissue Kit (Cat no. 69504, Valencia, CA). For the home and car sampling, swab was used for wiping the surface of cushion and carpet in car, and toy and toilet seat in home. Specifically, wipe the swab for 20 times on an area of 4×4cm ^2^. After collection, transfer the disposable swab to the tube containing sample storage liquid immediately, and break the swab along the crease. Then close the lid of the tube and make sure that there is no leakage. Finally, put the sample preservation solution tube in the biosafety bag for DNA extraction.

#### DNA extraction, 16S rRNA and shotgun metagenomic sequencing

Genomic DNA was extracted from each fecal containing tube using the Tissue and Blood DNA Isolation kit (Qiagen, Valencia, CA) following the manufacturer’s instructions with slight modifications. PCR amplification of the V1-V3 hypervariable regions of 16S rRNA genes was performed using the primer set (27F/534R) and followed the protocol developed by the Human Microbiome Project. PCR amplification reactions in triplicate for each sample were pooled at approximately equal amounts and sequenced, via the Illumina MiSeq 250 platform. All sequences were pre-processed following the standard QIIME (v.1.9) pipeline. Downstream bioinformatics analysis was performed using Parallel-Meta 3, a software package for comprehensive taxonomic and functional comparison of microbial communities. Clustering of OTUs was conducted at the 97% similarity level using a pre-clustered version of the Refseq database.

Paired-end metagenomic sequencing was performed for the two mock samples and fecal microbiota from three individuals via the Illumina HiSeq 2500 platform, yielding 239.73 ± 12.34 GB per sample (for fecal samples, with average fragment insert size of 350 bp and average read length of 150 bp). The reads were quality controlled by Trimmomatic (Sliding window 4:20; Minlength:100; MinPhred:25; Percentage of MinPhred: 80), and finally 858,032,764 ± 13,140,670 clean reads per sample were generated, and then profiled by mOTUs2 using default parameters.

#### 2bRAD-M sequencing

The 2bRAD-M library preparation basically followed the original protocol developed by Wang et al. (2012) ^15^ with minor modifications. Library preparation began with the digestion of 1pg–200 ng genomic DNA in a 15-µl reaction using 4 U BcgI (NEB) at 37 °C for 3 h. 5ul of digested product was run on a 1% agarose gel to verify digestion. Next, ligation reaction was performed at 4 °C for 16 h in a 20 µl volume containing 10 µl of di gested product, 0.2 µM each of library-specific adaptors (Ada1 and Ada2), 1 mM ATP (NEB), 1×T4 DNA Ligase Buffer and 800 U T4 DNA ligase (NEB). Then heat inactivation was performed for BcgI at 65 °C for 20 min.

Ligation products were amplified in 40-µl PCRs, each composed of 7 µl ligated DNA, 0.1 µM each primer (Primer1 and Primer2 for Illumina), 0.3 mM dNTP, 1× Phusion HF buffer and 0.4 U Phusion high fidelity DNA polymerase (NEB). PCR was conducted in a DNA Engine Tetrad 2 thermal cycler (Bio-Rad) with 16-28 cycles of 98 °C for 5 s, 60 °C for 20 s and 72 °C for 10 s and then a final extension of 10 min at 72 °C. The target band (Illumina: ~100 bp) was excised from 8% (wt/vol) polyacrylamide gel, and the DNA diffused from the gel into nuclease-free water for 12 h at 4 °C. Finally, barcodes were introduced by means of PCR with platform-specific barcode-bearing primers. 40-µl PCR reaction contained 50 ng of gel-extracted PCR product, 0.2 µM of each primer (Primer1 and Primer3 for Illumina), 0.6 mM dNTP, 1× Phusion HF buffer and 0.8 U Phusion high-fidelity DNA polymerase; seven cycles of the PCR profile listed above were performed. PCR products were purified by QIAquick PCR purification kit (Qiagen, Valencia, CA) and subjected to Illumina HiSeq platform sequencing. All primer and adaptor sequences are provided at **Table S8**.

### Data analysis

#### Identification of species-specific 2bRAD-M markers from the most comprehensive genome database

Firstly, totally 173,165 microbial genomes (including bacteria, fungi and archaea) were downloaded from NCBI RefSeq database. Then, built-in Perl scripts (GitHub: https://github.com/shihuang047/2bRAD-M) were used to sample restriction fragments from microbial genomes by each of 16 type 2B restriction enzymes, which formed a huge 2bRAD microbial genome database. The set of 2bRAD tags sampled from each genome was assigned under the GCF number, as well as GCF’s taxonomic information corresponding to the whole genome. Finally, all 2bRAD tags from each GCF that occur once within the genome were compared with those to all the others. Those 2bRAD tags are specific to a species-level taxon (having no overlap with other species’) were developed as species-specific 2bRAD markers, collectively forming a 2bRAD marker database.

#### Simulation of 2bRAD-M sequencing data

To test the generalizability of our 2bRAD markers for microbial profiling, we designed a microbiome structure containing 50 bacterial species from a wide range of habitats such as oral, gut and soil environments. “Wgsim” (https://github.com/lh3/wgsim) was then used (via default parameters) to simulate 150 bp long WMS reads according to specified species-level composition. A built-in script for digital restriction digestion was applied to simulate the 2bRAD-M sequencing data from the 150-bp WMS reads.

To benchmark 2bRAD-M with other metagenomic profilers, we further simulate WMS data of typical gut microbiomes (N=5). Firstly, we simulated gut microbiome profiles based on 100-150 species members that commonly identified in the real stool samples. Their abundances were created randomly from a log-normal distribution using “rlnorm” function in R with parameters: meanlog = 0 and sdlog = 1, and 10 repeats were simulated for the species number in each sample. Next, we also applied Wgsim to simulate the sequencing data given those fixed species compositions. Our 2bRAD-M should pre-process the WMS reads by transforming them into the 2bRAD format (e.g., 32-bp) as mentioned above.

#### Calculation of relative abundance

Firstly, to identify microbial species within each sample, all sequenced 2bRAD tags after quality control were mapped (using a built-in Perl script) against the 2bRAD marker database which contains all 2bRAD tags theoretically unique to each of 26,163 microbial species in RefSeq database (**Fig. 1**). To control the false-positive in the species identification, G score was derived for each species identified within a sample as below, which is a geometric mean of read coverage of 2bRAD markers belongs to a species and number of all detected 2bRAD markers of this species. The threshold of G score for a false positive discovery of microbial species was set as 10.

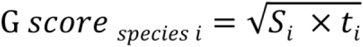

S: the number of reads assigned to all 2bRAD markers belonging to species *i* within a sample.
t: number of all 2bRAD markers of species *i* that have been sequenced within a sample

We estimated the relative abundance of each microbial species in a sample using the formula as below. We first calculated the average read coverage of all 2bRAD markers for each species, which represent the number of individuals belonging to a species present in a sample at a given sequencing depth. The relative abundance of a given species is then calculated as the ratio of the number of microbial individuals belonging to a species against the total number of individuals from known species that can be detected within a sample, with the default G score of 10.

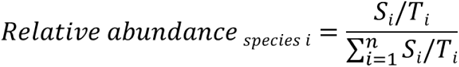

S: the number of reads assigned to all 2bRAD markers of species *i* within a sample
T: the number of all theoretical 2bRAD markers of species *i*

#### Calculation of Precision, Recall and L2 similarity

For overall performance assessment, we applied precision and recall to evaluate the accuracy of microbial identification, while L2 distance^16^ was employed to evaluate the accuracy of abundance estimation within a sample. Precision is the proportion of true positive species against the total number of species identified by a method, whereas recall is defined as the proportion of true positive species against the number of all species actually existed in a sample. To evaluate the accuracy of abundance profiles, we calculate the L2 distance between ground-truth abundance profile to each of taxonomic profiles at a given taxonomic level (e.g., species or genus) produced by metagenomic sequencing methods. To visualize the benchmarking results more intuitively, we further employed L2 similarity calculated as 1-L2 distance for performance comparisons.

#### Diagnosis model of cancer samples

Random forest models were trained to identify cancer status using the taxonomy profiles on the species level. Default parameters of the R implementation of algorithm were applied (R package ‘randomForest’, ntree=5,000, using default mtry of p/3 where p is the number of input taxa). The performance of RF models based on microbiota was evaluated with a ten-fold cross-validation approach.

Additional details are provided in **Supplemental Methods**.

## Acknowledgments

We thank Dr. Rob Knight and Dr. Yoshiki Vázquez-Baeza from UCSD for the helpful discussion on technical issues. This work was funded by Grant 31800088 from National Natural Science Foundation and 2019M652501 from China Postdoctoral Science Foundation. S. W. acknowledges the support by Taishan Scholar Fund of Shandong Province of China.

## Data availability

The sequencing data of 16S, WMS and 2bRAD-M for MOCK (MOCK_CAS and MOCK_MSA1002) and FFPE (only 2bRAD-M sequencing data) were submitted to figshare: https://doi.org/10.6084/m9.figshare.12272360.v6. The sequencing data of 16S and 2bRAD-M for typical low biomass samples (underarm, home and car) and fecal samples were submitted to https://doi.org/10.6084/m9.figshare.13514960.v1. The ultra-deep WMS sequencing were submitted to SRA under PRJNA689204 in NCBI.

## Code availability

The 2bRAD-M computational pipeline and related database files are publicly available at Github: https://github.com/shihuang047/2bRAD-M. All source data and codes for generation of figures and tables in the manuscript can be downloaded from: https://github.com/shihuang047/2bRAD-M-manuscript.

## Author contributions

Z.S. and S.H. conceived of the study. S.W. and J.X. designed the experiments and supervised the whole study. P.Z., J.L., L.Z, Q.N. and X.W carried out the experiments. Z.S., S.H., S.W. and J.X. wrote the manuscript with support from L.T., H.Z., Z.B. and J.L.. Z.S., G.J. and M.Z. planned and carried out the simulations. Z.S. and R.Z. designed the model and the computational framework and analyzed the data.

