## Supplemental information for "Species-resolved sequencing of low-biomass microbiomes by 2bRAD-M"

**Supplemental Methods**

**Supplemental Tables and Figures**

**Table S1.** Availability of 2bRAD-M markers for taxonomic profiling at each of the taxonomic levels.

**Table S2.** Expected abundance of bacterial species in simulation data and profiling results by the 2bRAD-M computational pipeline.

**Table S3.** The relative abundance of major taxa identified in the three fecal samples at the species level using 2bRAD-M or WMS or at the genus level using 16S rRNA gene amplicon sequencing.

**Table S4.** The initial DNA content and metadata for underarm skin, home and car samples.

**Table S5.** The 2bRAD-M profiling results for underarm, home and car samples.

**Table S6.** The relative abundance of bacteria, fungi and archaea in the indoor built-environmental samples.

**Table S7.** Species-level microbial organismal markers for the highly reliable diagnosis of cervical cancer from FFPE samples.

**Table S8.** The adaptors and primers used in 2bRAD-M sequencing (5'-3').

**Figure S1.** The theoretical fragments generated by 2bRAD-M and their originated genomes.

**Figure S2.** Rarefaction analysis reveals the desired sequencing depth for 2bRAD-M and WMS for reliable taxonomic profiling.

**Figure S3.** Comparison of the genus-level taxonomic profiles based on 16S rRNA sequencing and 2bRAD-M in each of the underarm, home or car samples.

**Figure S4.** Rarefaction analysis reveals the desired sequencing depth of 2bRAD-M for taxonomic profiling of the representative built-environment and FFPE samples.

**Figure S5**. Species abundance profiles of the FFPE samples from healthy tissue, pre-invasive cancer and invasive cancer.

**Figure S6.** Agarose gel analysis of the DNA extracted from cervical FFPE tissue samples.

### Supplementary Methods

#### Feasibility of 2bRAD-M for microbiome profiling by additional type IIB restriction enzymes

To test the feasibility of 2bRAD-M for microbiome profiling, two fundamental questions would be addressed in the *in silico* experiments based on an extensive set of microbial genomes: (*i*) whether the surveyed 2bRAD-M data is a reliable reduced representation of the microbial genomes; (*ii*) whether the surveyed 2bRAD-M data harbor phylogenetic markers that can enable the taxonomic profiling of microbial taxa at the species level.

We started by downloading 173,165 microbial genomes from NCBI RefSeq (Oct, 2019), including 15,162 bacterial, archaea and fungal complete genomes. The digital restriction digestion of all these genomes by an IIB restriction enzyme (such as BcgI) resulted in averagely 2930.38±2790.84 2bRAD-M tags per genome. To date, there are totally 16 type IIB restriction enzymes discovered, and they have distinct DNA recognition sites. We thus performed the digital digestion of all microbial genomes using all these 16 type IIB restriction enzymes, which produced multiple and flexible reduced representations of each microbial genome. Collectively, we identified and collected the restriction fragments of all microbial genomes using 16 restriction enzymes, which represent the most comprehensive 2bRAD-M reference genome database.

To assess whether the surveyed restriction fragments represent a random subset of a given microbial genome, we compared a number of features of the digitally digested DNA fragments from a given microbial genome to those from the entire genome. We found that the surveyed fragments are typically evenly distributed along a microbial genome, and across genic and non-genic regions (**Fig S1**). Likewise, %G + C content (53%) of surveyed 2bRAD-M tags are very similar to the genome-wide averages (Pearson's correlation R=0.992). Furthermore, the number of

2bRAD-M tags is highly correlated with the genome size of a given microbe (Pearson's correlation R=0.976). This suggests that 2bRAD-M fragments can be employed to survey genome-wide features of microbes without requiring sequencing the full genome, regardless of the specific type-2B enzyme used here (**Fig. S1**).

We next sought to identify universal 2bRAD-M phylogenetic markers from a total set of 173,165 reference microbial genomes. Different strategies have been introduced to determine microbial community compositions and estimate their abundances from metagenomic data. Our approach is to identify taxa-specific DNA markers by analyzing the 2bRAD-M reference genome database and further quantify the read coverage of those markers for taxonomic profiling from 2bRAD-M data. Therefore, desired DNA markers in our study should be specific to taxa (i.e., species), iso-length (around 33 bp long) and short DNA fragments that only occur once per genome. Overall, the higher taxonomic level, the more 2bRAD-M tags are available. At the Kingdom level, almost all 2bRAD-M tags are kingdom-specific, thus there are very few shared 2bRAD-M tags among bacteria, fungi, archaea and human, regardless of the restriction enzymes. This suggested that the abundance ratio between the kingdoms can be readily derived from the 2bRAD-M data (yet can be challenging for WMS [1]). The phylum-specific 2bRAD-M markers accounted for up to 90% ~ 97% of all theoretical 2bRAD-M tags produced from a given restriction enzyme from a given microbial genome. We next explored the 2bRAD-M markers specific to the 26,163 microbial species. Among all 521,289,189 restriction fragments produced by one typical type IIB restriction enzyme (BcgI), 99.21% are single-copy within a given microbial genome, while averagely 21.86% are specific to species-level taxa (**Table S1**). The other restriction enzymes can also generate distinct sets of 2bRAD-M tags from each microbial genome. In fact,

18.81%-25.80% single-copy species-specific markers were identified from the 2bRAD-M genomes digested by the other restriction enzymes. Therefore, in principle, 2bRAD-M data provides a rich and highly flexible source of phylogenetic markers for metagenomic profiling.

**The benchmarking of taxonomic profiling methods on real fecal samples**

To profile the fecal microbiomes, three sequencing approaches were used for each sample. (*i*) Deep WMS generated averagely 429,016,382 clean reads per sample (about 239.73 GB per sample) by HiSeq 2500. Then mOTUs2 was employed with default parameters and RefSeq database to identify the microbial compositions. (*ii*) Partial 16S rRNA gene amplicon (V1-V3) sequencing generated averagely 100,340 clean reads per sample (about 0.084 GB per sample) by Miseq 250. The merged clean reads were then searched against RefSeq for classification. (*iii*) 2bRAD-M sequencing generates averagely 1,234,122 clean tags (about 0.254 GB per sample) using Illumina HiSeq X Ten platform. Then these high-quality 2bRAD-M tags were mapped against the 2bRAD-M marker database for the species-level taxa. As 16S rRNA sequencing was involved here, bioinformatic tools using the marker-gene-based strategy for taxonomic assignment should be adopted by the 2bRAD-M and WMS methods to profile those samples. Therefore, we employed the Parallel-Meta3 pipeline[2] on 16S rRNA sequencing data, the 2bRAD-M bioinformatic pipeline (from this study) on 2bRAD-M data, and mOTU[3] on the WMS data respectively, since all these methods are based on marker gene databases.

### Supplementary Tables

**Table S1. Availability of 2bRAD-M markers for taxonomic profiling at each of the taxonomic levels.** In each row, the value indicates the average percentage of taxa-specific 2bRAD-M tags in all 2bRAD-M tags produced by a given type IIB enzyme. Those 2bRAD-M marker tags are all single-copy in a microbial genome and specific to a given taxon. Thus for each of the type IIB enzymes and at each of the taxonomic levels, 2bRAD-M markers or taxonomic profiling are abundant.

| IIB enzyme | Phylum | Class | Order | Family | Genus | Species |
|---|---|---|---|---|---|---|
| **AlfI** | 89.12% | 86.98% | 82.84% | 79.44% | 72.94% | 39.79% |
| **AloI** | 86.81% | 84.30% | 79.41% | 75.37% | 68.75% | 36.85% |
| **BaeI** | 87.50% | 85.35% | 81.16% | 77.76% | 71.53% | 39.19% |
| **BcgI** | 88.85% | 86.64% | 82.54% | 79.23% | 72.71% | 39.68% |
| **BplI** | 87.14% | 84.58% | 80.57% | 76.96% | 70.17% | 38.16% |
| **BsaXI** | 87.39% | 85.08% | 80.62% | 77.03% | 70.61% | 38.62% |
| **BslFI** | 86.17% | 83.96% | 79.65% | 76.30% | 69.95% | 37.90% |
| **Bsp24I** | 87.16% | 84.88% | 80.41% | 76.85% | 70.22% | 38.10% |
| **CjeI** | 87.58% | 85.35% | 80.99% | 77.46% | 70.89% | 38.57% |
| **CjePI** | 87.85% | 85.56% | 81.15% | 77.51% | 70.94% | 38.29% |
| **CspCI** | 88.88% | 86.68% | 82.95% | 80.09% | 74.09% | 41.89% |
| **FalI** | 86.77% | 84.24% | 79.21% | 75.37% | 68.45% | 37.08% |
| **HaeIV** | 87.43% | 85.17% | 80.89% | 77.35% | 70.93% | 38.56% |
| **Hin4I** | 86.98% | 84.79% | 80.52% | 77.01% | 70.64% | 38.56% |
| **PpiI** | 87.93% | 85.24% | 80.56% | 76.55% | 69.97% | 37.33% |
| **PsrI** | 84.96% | 82.43% | 77.53% | 73.60% | 66.78% | 36.28% |

**Table S2. Expected abundance of bacterial species in simulation data and profiling results from the 2bRAD-M computational pipeline.**

| Organism Name | Assembly Accession | Relative abundance | 2bRAD-M |
|---|---|---|---|
| *Archaeoglobus fulgidus* DSM 4304 | GCF_000008665.1 | 0.667% | 0.673% |
| *Clostridium acetobutylicum* ATCC 824 | GCF_000008765.1 | 0.667% | 0.000% |
| *Lactobacillus salivarius* UCC118 | GCF_000008925.1 | 0.667% | 0.716% |
| *Ralstonia solanacearum* GMI1000 | GCF_000009125.1 | 0.667% | 0.663% |
| *Nitrosomonas europaea* ATCC 19718 | GCF_000009145.1 | 0.667% | 0.697% |
| *Helicobacter acinonychis str.* Sheeba | GCF_000009305.1 | 0.667% | 0.704% |
| *Yersinia enterocolitica subsp. enterocolitica* 8081 | GCF_000009345.1 | 0.667% | 0.659% |
| *Alcanivorax borkumensis* SK2 | GCF_000009365.1 | 0.667% | 0.657% |
| *Streptococcus uberis* 0140J | GCF_000009545.1 | 0.667% | 0.648% |
| *Staphylococcus haemolyticus* JCSC1435 | GCF_000009865.1 | 0.667% | 0.758% |
| *Symbiobacterium thermophilum IAM 14863* | GCF_000009905.1 | 1.333% | 1.370% |
| *Chlamydia felis* Fe/C-56 | GCF_000009945.1 | 1.333% | 1.315% |
| *Thermococcus kodakarensis* KOD1 | GCF_000009965.1 | 1.333% | 1.368% |
| *Magnetospirillum magneticum* AMB-1 | GCF_000009985.1 | 1.333% | 1.372% |
| *Synechococcus elongatus* PCC 6301 | GCF_000010065.1 | 1.333% | 1.230% |
| *Sodalis glossinidius str. 'morsitans'* | GCF_000010085.1 | 1.333% | 1.311% |
| *Finegoldia magna* ATCC 29328 | GCF_000010185.1 | 1.333% | 1.428% |
| *Gemmatimonas aurantiaca* T-27 | GCF_000010305.1 | 1.333% | 1.317% |
| *Nitratiruptor sp.* SB155-2 | GCF_000010325.1 | 1.333% | 1.298% |
| *Sulfurovum sp.* NBC37-1 | GCF_000010345.1 | 1.333% | 1.330% |
| *Bifidobacterium adolescentis* ATCC 15703 | GCF_000010425.1 | 2.000% | 1.908% |
| *Porphyromonas gingivalis* ATCC 33277 | GCF_000010505.1 | 2.000% | 1.981% |
| *Azorhizobium caulinodans* ORS 571 | GCF_000010525.1 | 2.000% | 2.018% |
| *Macrococcus caseolyticus* JCSC5402 | GCF_000010585.1 | 2.000% | 2.076% |
| *Candidatus Azobacteroides pseudotrichonymphae genomovar. CFP2* | GCF_000010645.1 | 2.000% | 2.590% |
| *Acetobacter pasteurianus IFO* 3283-01 | GCF_000010825.1 | 2.000% | 2.048% |
| *Deferribacter desulfuricans* SSM1 | GCF_000010985.1 | 2.000% | 2.011% |
| *Pyrococcus horikoshii* OT3 | GCF_000011105.1 | 2.000% | 2.058% |
| *Thermoplasma volcanium* GSS1 | GCF_000011185.1 | 2.000% | 1.996% |
| *Mycoplasma penetrans* HF-2 | GCF_000011225.1 | 2.000% | 2.035% |
| *Oceanobacillus iheyensis* HTE831 | GCF_000011245.1 | 2.667% | 2.651% |
| *Thermosynechococcus elongatus* BP-1 | GCF_000011345.1 | 2.667% | 2.695% |
| *Gloeobacter violaceus* PCC 7421 | GCF_000011385.1 | 2.667% | 2.657% |
| *Ruegeria pomeroyi* DSS-3 | GCF_000011965.2 | 2.667% | 2.626% |
| *Rickettsia felis* URRWXCal2 | GCF_000012145.1 | 2.667% | 2.720% |
| *Psychrobacter arcticus* 273-4 | GCF_000012305.1 | 2.667% | 2.640% |

| | | | |
|---|---|---|---|
| *Thermobifida fusca* YX | GCF_000012405.1 | 2.667% | 2.803% |
| *Dechloromonas aromatica* RCB | GCF_000012425.1 | 2.667% | 2.641% |
| *Pelodictyon luteolum* DSM 273 | GCF_000012485.1 | 2.667% | 2.627% |
| *Synechococcus sp.* CC9902 | GCF_000012505.1 | 2.667% | 2.642% |
| *Methanosphaera stadtmanae* DSM 3091 | GCF_000012545.1 | 3.333% | 3.357% |
| *Ehrlichia canis str.* Jake | GCF_000012565.1 | 3.333% | 3.232% |
| *Chlorobium chlorochromatii* CaD3 | GCF_000012585.1 | 3.333% | 3.308% |
| *Nitrobacter winogradskyi* Nb-255 | GCF_000012725.1 | 3.333% | 3.073% |
| *Nitrosococcus oceani* ATCC 19707 | GCF_000012805.1 | 3.333% | 3.387% |
| *Carboxydothermus hydrogenoformans* Z-2901 | GCF_000012865.1 | 3.333% | 3.351% |
| *Pelobacter carbinolicus* DSM 2380 | GCF_000012885.1 | 3.333% | 3.375% |
| *Sulfurimonas denitrificans* DSM 1251 | GCF_000012965.1 | 3.333% | 3.292% |
| *Alternaria arborescens* | GCF_004154835.1 | 3.333% | 3.334% |
| *Apiotrichum porosum* | GCF_003942205.1 | 3.333% | 3.345% |
| *Alternaria_alternata* | GCF_001642055.1 | 0.000% | 0.006% |

**Table S3. The relative abundance of major taxa identified in the three fecal samples at the species level using 2bRAD-M or WMS or at the genus level using 16S rRNA gene amplicon sequencing.**

| Real fecal sample A | | | | | | | |
|---|---|---|---|---|---|---|---|
| **Top species in 2bRAD-M** | **Relative abundance** | **Rank** | **Corresponding relative abundance in WMS** | **Rank** | **Corresponding genus in 16S** | **Relative abundance** | **Rank** |
| *Prevotella copri* | 61.86% | 1 | 67.30% | 1 | *Prevotella* | 71.14% | 1 |
| *Prevotella sp BCRC* | 10.81% | 2 | NA | NA | *Prevotella* | 71.14% | 1 |
| *Prevotella stercorea* | 4.85% | 3 | 9.07% | 2 | *Prevotella* | 71.14% | 1 |
| *Bacteroides plebeius* | 2.97% | 4 | 3.20% | 3 | *Bacteroides* | 16.44% | 2 |
| *Bacteroides coprophilus* | 1.46% | 5 | 1.68% | 5 | *Bacteroides* | 16.44% | |
| *Bacteroides uniformis* | 1.37% | 6 | NA | NA | *Bacteroides* | 16.44% | |
| *Bacteroides coprocola* | 1.31% | 7 | 1.43% | 7 | *Bacteroides* | 16.44% | |
| *Bacteroides thetaiotaomicron* | 0.95% | 8 | NA | NA | *Bacteroides* | 16.44% | |
| *Acinetobacter baumannii* | 0.81% | 9 | NA | NA | *NA* | NA | NA |
| *Bacteroides stercoris* | 0.71% | 10 | 0.63% | 14 | *Bacteroides* | 16.44% | 2 |
| *Alistipes putredinis* | 0.69% | 11 | 1.35% | 8 | *Alistipes* | 0.89% | 6 |
| *Prevotella sp AM23 5* | 0.65% | 12 | NA | NA | *Prevotella* | 71.14% | 1 |
| *Parabacteroides merdae* | 0.61% | 13 | 0.64% | 13 | *Parabacteroides* | 1.93% | 4 |
| *Parabacteroides distasonis* | 0.59% | 14 | 0.51% | 16 | *Parabacteroides* | 1.93% | 4 |
| *Bacteroides massiliensis* | 0.57% | 15 | 0.74% | 11 | *Bacteroides* | 16.44% | 2 |
| *Bacteroides vulgatus* | 0.47% | 16 | NA | NA | *Bacteroides* | 16.44% | 2 |
| *Eubacterium rectale* | 0.41% | 17 | 0.41% | 178 | *NA* | NA | NA |
| *Megamonas funiformis* | 0.38% | 18 | NA | NA | *Megamonas* | 0.71% | 9 |
| *Phascolarctobacterium succinatutens* | 0.33% | 19 | 0.53% | 14 | *Phascolarctobacterium* | 0.56% | 11 |
| *Bacteroides salyersiae* | 0.31% | 20 | 0.31% | 21 | *Bacteroides* | 16.44% | 2 |
| **SUM** | 92.10% | | 87.79% | | | 91.67% | |

| Real fecal sample B | | | | | | | |
|---|---|---|---|---|---|---|---|
| **Top species in 2bRAD-M** | **Relative abundance** | **Rank** | **Corresponding relative abundance in WMS** | **Rank** | **Corresponding genus in 16S** | **Relative abundance** | **Rank** |
| *Prevotella copri* | 58.00% | 1 | 58.76% | 1 | *Prevotella* | 69.47% | 1 |
| *Prevotella sp BCRC* | 7.23% | 2 | NA | NA | *Prevotella* | 69.47% | 1 |
| *Prevotella stercorea* | 4.36% | 3 | 8.82% | 2 | *Prevotella* | 69.47% | 1 |
| *Bacteroides coprophilus* | 4.26% | 4 | 4.70% | 3 | *Bacteroides* | 13.49% | 2 |
| *Acinetobacter baumannii* | 3.13% | 5 | NA | NA | *NA* | NA | NA |

| | | | | | | | |
|---|---|---|---|---|---|---|---|
| *Eubacterium rectale* | 2.83% | 6 | 3.01% | 4 | *NA* | NA | NA |
| *Bacteroides plebeius* | 1.72% | 7 | 1.85% | 8 | *Bacteroides* | 13.49% | 2 |
| *Bacteroides vulgatus* | 1.59% | 8 | NA | NA | *Bacteroides* | 13.49% | 2 |
| *Bacteroides dorei* | 1.16% | 9 | NA | NA | *Bacteroides* | 13.49% | 2 |
| *Bacteroides uniformis* | 1.10% | 10 | NA | NA | *Bacteroides* | 13.49% | 2 |
| *Parabacteroides merdae* | 1.00% | 11 | 1.39% | 9 | *Parabacteroides* | 1.86% | 4 |
| *Alistipes putredinis* | 0.93% | 12 | 2.33% | 5 | *Alistipes* | 0.81% | 9 |
| *Lachnospira pectinoschiza* | 0.71% | 13 | NA | NA | *NA* | NA | NA |
| *Sutterella sp KLE1602* | 0.64% | 14 | NA | NA | *Sutterella* | 1.26% | 7 |
| *Bacteroides stercoris* | 0.58% | 15 | 0.62% | 14 | *Bacteroides* | 13.49% | 2 |
| *Parabacteroides distasonis* | 0.56% | 16 | 0.62% | 15 | *Parabacteroides* | 1.86% | 4 |
| *Prevotella sp Marseille* | 0.51% | 17 | NA | NA | *Prevotella* | 69.47% | 1 |
| *Bacteroides caccae* | 0.49% | 18 | 0.46% | 19 | *Bacteroides* | 13.49% | 2 |
| *Dialister sp Marseille* | 0.46% | 19 | NA | NA | *Dialister* | 0.01% | 36 |
| *Megamonas funiformis* | 0.43% | 20 | NA | NA | *Megamonas* | 1.70% | 5 |
| **SUM** | 91.67% | | 82.56% | | | 88.60% | |
| **Real fecal sample C** | | | | | | | |
| **Top species in 2bRAD-M** | **Relative abundance** | **Rank** | **Corresponding relative abundance in WMS** | **Rank** | **Corresponding genus in 16S** | **Relative abundance** | **Rank** |
| *Prevotella copri* | 29.07% | 1 | 33.40% | 1 | *Prevotella* | 29.30% | 1 |
| *Bacteroides massiliensis* | 8.35% | 2 | 8.71% | 2 | *Bacteroides* | 24.98% | 2 |
| *Bacteroides dorei* | 6.28% | 3 | NA | NA | *Bacteroides* | 24.98% | 2 |
| *Faecalibacterium prausnitzii* | 5.75% | 4 | 7.61% | 3 | *Faecalibacterium* | 11.65% | 3 |
| *Bacteroides plebeius* | 5.17% | 5 | 5.52% | 5 | *Bacteroides* | 24.98% | 2 |
| *Bacteroides uniformis* | 3.78% | 6 | NA | NA | *Bacteroides* | 24.98% | 2 |
| *Roseburia intestinalis* | 3.29% | 7 | 2.41% | 10 | *Roseburia* | 8.48% | 4 |
| *Alistipes putredinis* | 3.03% | 8 | 5.82% | 4 | *Alistipes* | 2.05% | 8 |
| *Roseburia inulinivorans* | 2.95% | 9 | 1.79% | 11 | *Roseburia* | 8.48% | 4 |
| *Lachnospira pectinoschiza* | 2.43% | 10 | NA | NA | *NA* | NA | NA |
| *Clostridium sp AM42* | 1.35% | 11 | NA | NA | *Clostridium* | 3.12% | 7 |
| *Clostridium sp AF43* | 1.23% | 12 | NA | NA | *Clostridium* | 3.12% | 7 |
| *Bacteroides caccae* | 1.13% | 13 | 0.89% | 19 | *Bacteroides* | 24.98% | 2 |
| *Roseburia faecis* | 1.11% | 14 | NA | NA | *Roseburia* | 8.48% | 4 |
| *Bacteroides sp* | 1.08% | 15 | NA | NA | *Bacteroides* | 24.98% | 2 |
| *Eubacterium rectale* | 1.03% | 16 | 0.97% | 18 | *NA* | NA | NA |
| *Firmicutes bacterium OM08* | 1.02% | 17 | NA | NA | *NA* | NA | NA |
| *Paraprevotella clara* | 1.00% | 18 | 1.59% | 12 | *Paraprevotella* | 1.41% | 11 |
| *Firmicutes bacterium AF22* | 0.96% | 19 | NA | NA | *NA* | NA | NA |
| *Clostridium sp OM08* | 0.93% | 20 | NA | NA | *Clostridium* | 3.12% | 7 |
| **SUM** | 80.94% | | 68.70% | | | 80.97% | |

**Table S4. The initial DNA content and metadata for underarm skin, home and car samples.**

| Sample ID | Gender | Age | Ethnicity | Sampling Site | Concentration ng/μL(Qubit) |
|---|---|---|---|---|---|
| **UA_01** | Female | 26-35 | Chinese | Underarm | 0.5414 |
| **UA_02** | Female | 46-55 | Indian | Underarm | 0.09839 |
| **UA_03** | Male | 26-35 | Chinese | Underarm | 0.06578 |
| **UA_04** | Female | 46-55 | Chinese | Underarm | 1.489 |
| **UA_05** | Male | 26-35 | Indian | Underarm | 2.421 |
| **UA_06** | Male | 26-35 | Filipino | Underarm | 3.924 |
| **UA_07** | Female | 36-45 | Indian | Underarm | 0.5529 |
| **UA_08** | Male | 26-35 | Chinese | Underarm | 4.182 |
| **UA_09** | Female | 26-35 | Indian | Underarm | 24.49 |
| **UA_10** | Male | 36-45 | Chinese | Underarm | 1.535 |
| **UA_11** | Female | 36-45 | Chinese | Underarm | 1.472 |
| **UA_12** | Male | 26-35 | Chinese | Underarm | 0.1819 |
| **UA_13** | Female | 26-35 | Filipino | Underarm | 4.339 |
| **UA_14** | Male | 18-25 | Indian | Underarm | 2.096 |
| **UA_15** | Female | 46-55 | Filipino | Underarm | 4.911 |
| **UA_16** | Male | 46-55 | Filipino | Underarm | 3.938 |
| **UA_17** | Male | 46-55 | Filipino | Underarm | 1.793 |
| **UA_18** | Male | 26-35 | Chinese | Underarm | 1.214 |
| **UA_19** | Male | 36-45 | Indian | Underarm | 1.284 |
| **UA_20** | Male | 36-45 | Indian | Underarm | 1.379 |
| **Car_KC05T3C-1** | NA | NA | NA | Cushion in car | 10.99 |
| **Car_KC06BgP-1** | NA | NA | NA | Floor mat in car | 20.28 |
| **Car_KC06T1P-1** | NA | NA | NA | Floor mat in car | 35.84 |
| **Car_KC06T2P-1** | NA | NA | NA | Floor mat in car | 9.04 |
| **Car_KC06T3P-1** | NA | NA | NA | Floor mat in car | 11.83 |
| **Car_KCT0P-1** | NA | NA | NA | Floor mat in car | 19.22 |
| **Car_KT05BgC-1** | NA | NA | NA | Cushion in car | 11.96 |
| **Car_KT05T2C-1** | NA | NA | NA | Cushion in car | 10.43 |
| **Home_SMM-2-3** | NA | NA | NA | Child's book | 5.231 |
| **Home_SY-41** | NA | NA | NA | Toilet | 13.49 |
| **Home_WJM-I** | NA | NA | NA | Child's toy | 9.917 |
| **Home_WX-9** | NA | NA | NA | Toilet mat | 21.77 |

**Table S5. The 2bRAD-M profiling results for underarm, home and car samples (in a separate file).**

**Table S6. The relative abundance of bacteria, fungi and archaea in the indoor built-environmental samples.**

| Kingdom | Underarm | Home | Car |
|---|---|---|---|
| **Bacteria** | 99.69% | 99.93% | 98.71% |
| **Fungi** | 0.31% | 0.07% | 1.26% |
| **Archaea** | 0.00% | 0.001% | 0.03% |

**Table S7. Species-level microbial organismal markers for the highly reliable diagnosis of cervical cancer from FFPE samples.**

| Species name | Mean abundance | | | Importance |
|---|---|---|---|---|
| | Severer Cancer | Cancer | Health | |
| *Porphyrobacter cryptus* | 0.297% | 0.000% | 0.002% | 0.200911002 |
| *Pelomonas puraquae* | 1.487% | 0.437% | 0.039% | 0.057330947 |
| *Methyloversatilis discipulorum* | 13.050% | 2.573% | 0.547% | 0.056879324 |
| *Methyloversatilis universalis* | 0.024% | 0.011% | 0.016% | 0.052514646 |
| *Pseudomonas aeruginosa* | 1.161% | 0.233% | 0.039% | 0.029690798 |
| *Mycobacterium tuberculosis* | 25.149% | 28.064% | 7.960% | 0.02625783 |
| *Escherichia coli* | 31.957% | 33.057% | 62.531% | 0.018157477 |
| *Lactobacillus paracasei* | 1.251% | 1.748% | 0.686% | 0.012936985 |
| *Ferrovibrio sp K5* | 1.187% | 0.350% | 0.129% | 0.010616483 |

**Table S8. The adaptors and primers used in 2bRAD-M sequencing (5’-3’).**

| Name | Adaptor sequence |
|---|---|
| **Adap-1 sense** | ACACTCTTTCCCTACACGACGCTCTTCCGATCTNN |
| **Adap-2 sense** | GTGACTGGAGTTCAGACGTGTGCTCTTCCGATCTNN |
| **Adap antisense** | AGATCGGAAGAGC |
| | **Primer sequence** |
| **Primer1** | ACACTCTTTCCCTACACGACGCT |
| **Primer2** | GTGACTGGAGTTCAGACGTGTGCT |
| **Primer3** | AATGATACGGCGACCACCGAGATCTACACTCTTTCCCTACACGACGCT |
| **Index primer** | CAAGCAGAAGACGGCATACGAGATXXXXXXGTGACTGGAGTTCAGACGTGT |

### Supplementary Figures

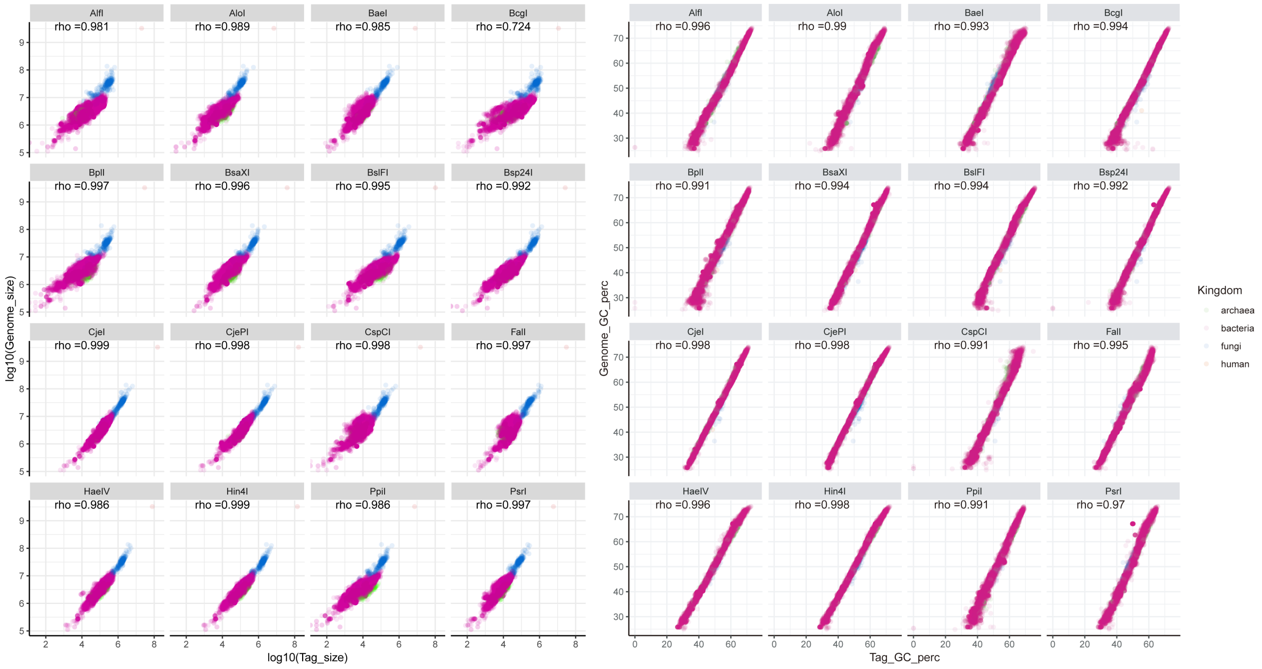


**Figure S1. The theoretical fragments generated by 2bRAD-M and their originated genomes.** Correlation of fragment size (left panel) or GC content (right panel) is shown. For a given genome, che collective size of all DNA tags cleaved by a type IIB restriction enzyme corresponds to a reduction in sequencing for one to two orders of magnitude (depending on genome size).

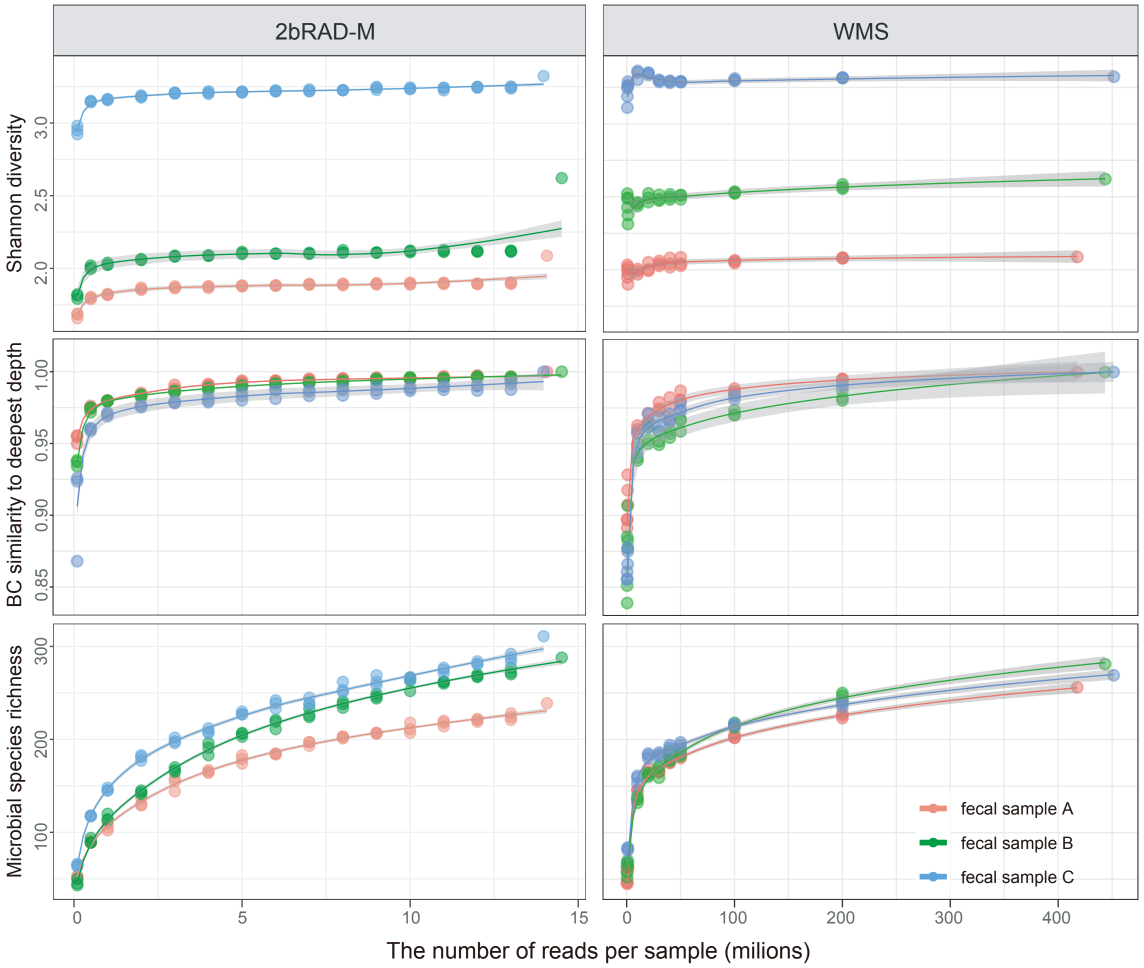


**Figure S2. Rarefaction analysis reveals the desired sequencing depth for 2bRAD-M and WMS for reliable taxonomic profiling.** In each scatter plot, we compared the profiling results of a fecal sample based on a method (either 2bRAD-M or WMS) at deep or shallow (by subsampling) depth of sequencing, via Shannon diversity, beta diversity (on the Bray-Curtis distance metric),and species richness. Based on the Shannon diversity and Bray-Curtis distance metrics, profiling results of 2bRAD-M quickly saturated at a shallow sequencing depth (2-3 million reads per sample). In contrast, for the same metrics, WMS-based taxonomic profiles saturated at far deeper sequencing depth (20-40 million reads per sample), suggesting much higher sequencing costs. The microbial richness (number of species-level taxa detected) based on both methods still grows as the sequencing depth increases, which is consistent with current knowledge on metagenomic diversity analysis.

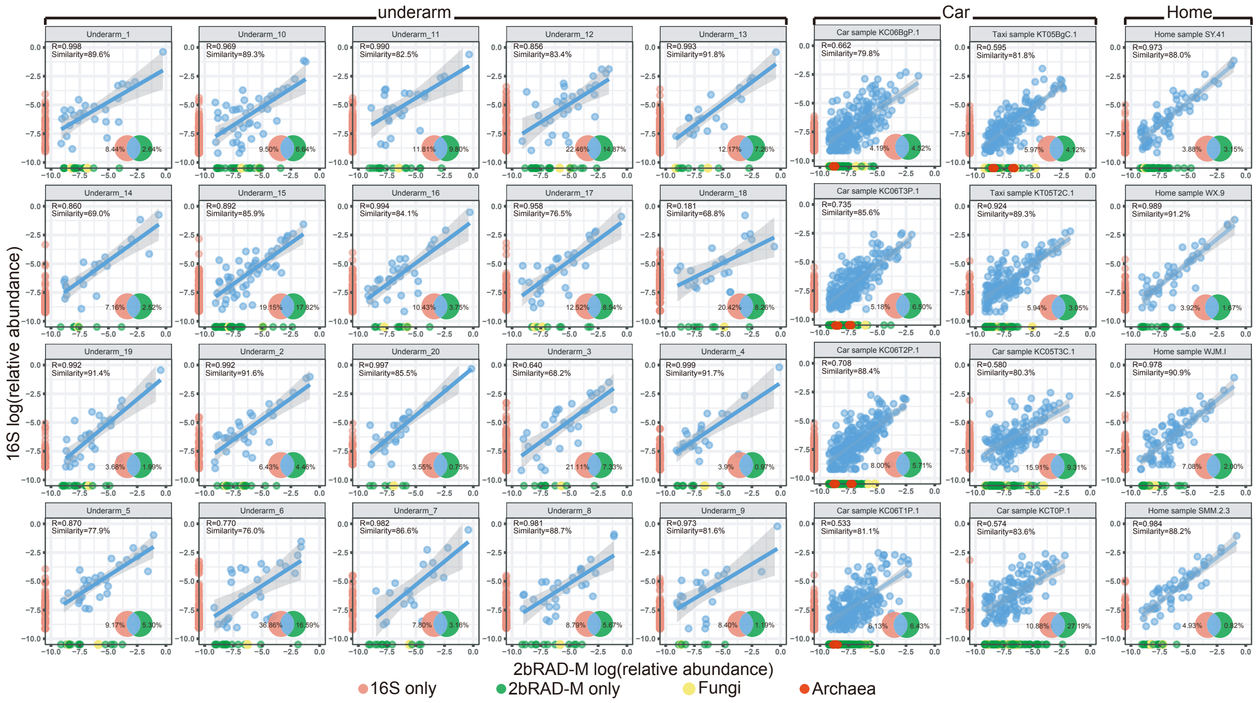


**Figure S3. Comparison of the genus-level taxonomic profiles based on 16S rRNA sequencing and 2bRAD-M in each of the underarm, home or car samples.** In each scatter plot, blue points represent the genus-level taxa shared between 16S rRNA sequencing and 2bRAD-M, while red points and green points refer to the unique genera identified by 16S rRNA sequencing or 2bRAD-M separately. Each yellow point represents a fungal taxon detected in a given sample by 2bRAD-M. The inset (Venn diagram) shows the overlapping fraction of identified taxa between 16S rRNA and 2bRAD-M profiles. Results for each of the 32 microbiome samples from underarm, car surfaces and home surfaces were presented.

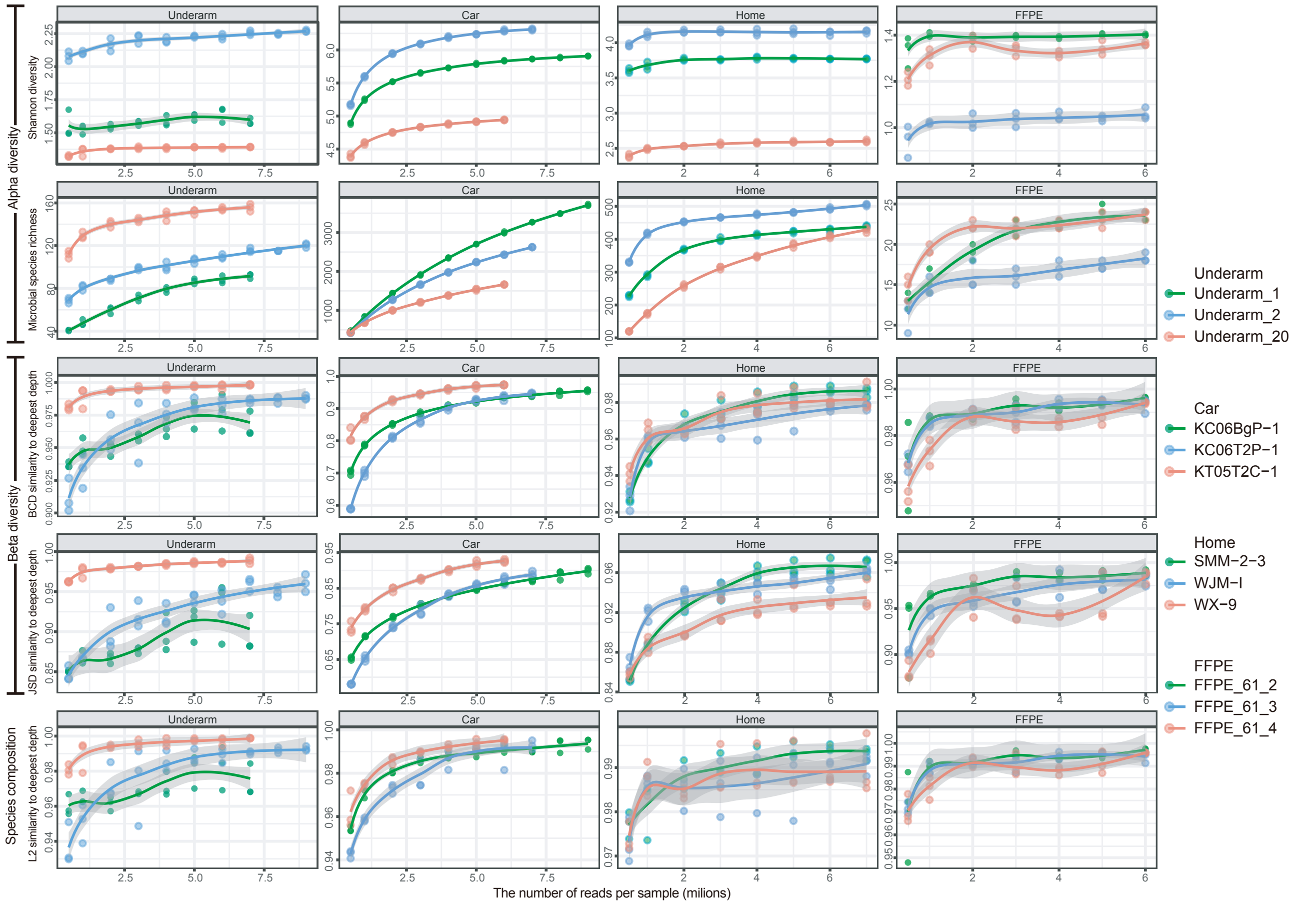


**Figure S4. Rarefaction analysis reveals the desired sequencing depth of 2bRAD-M for taxonomic profiling of the representative built-environment and FFPE samples.** From each of the categories of underarm skin, car surfaces, home surfaces and FFPE, three representative samples were shown as examples. For each sample, we compared key parameters of the taxonomic profiling (i.e., alpha diversity, beta diversity and species-level compositions) at several shallow sequencing depths (by subsampling) with those at the deep sequencing depth. The scatter plot in each panel indicates the relationship between a parameter ($y$-axis) and the sequencing depth ($x$-axis).

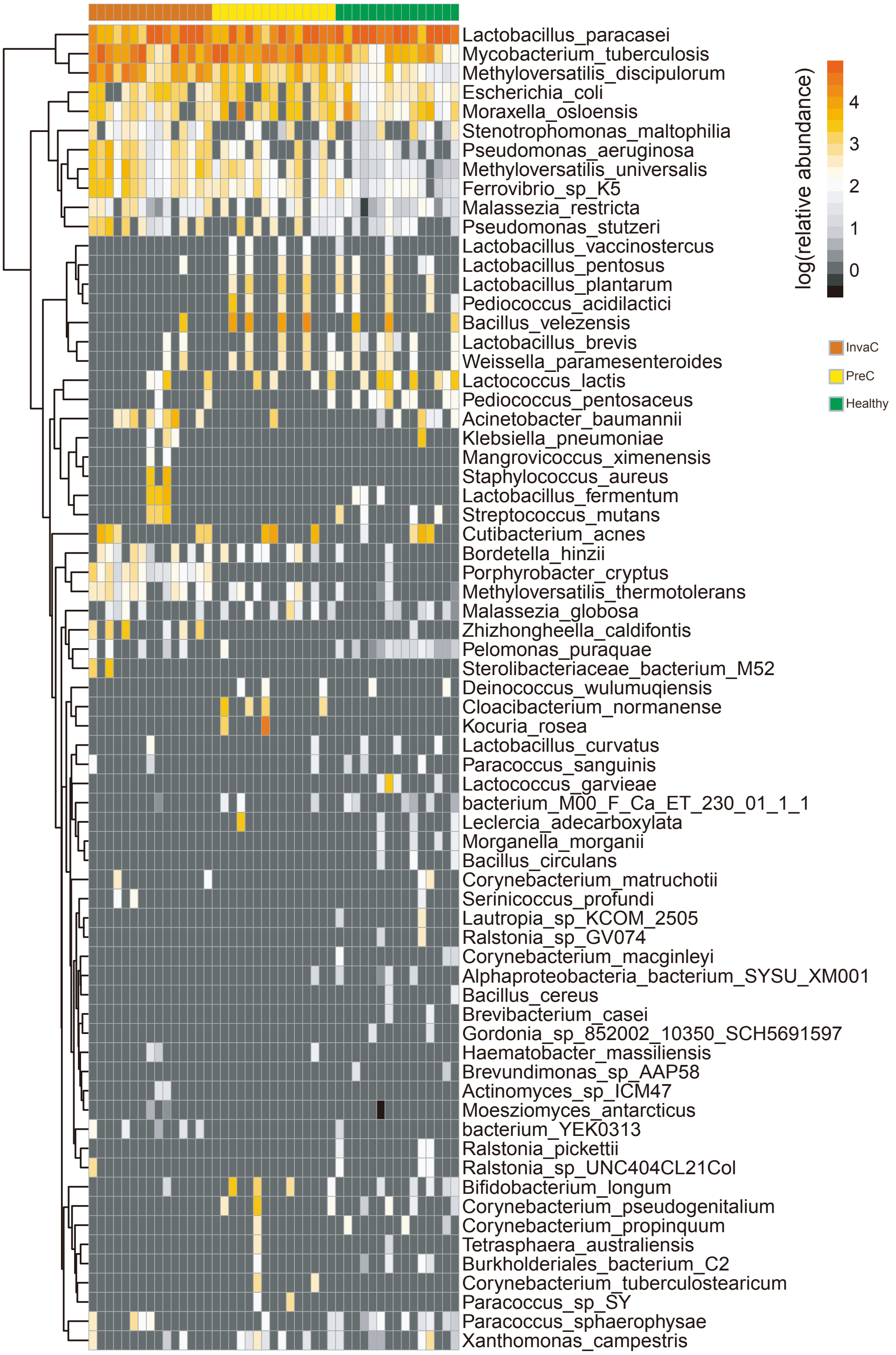


**Figure S5. Species abundance profiles of the FFPE samples from healthy tissue, pre-invasive cancer and invasive cancer.** The species with a positive importance score in the RF model were presented in the heat map.

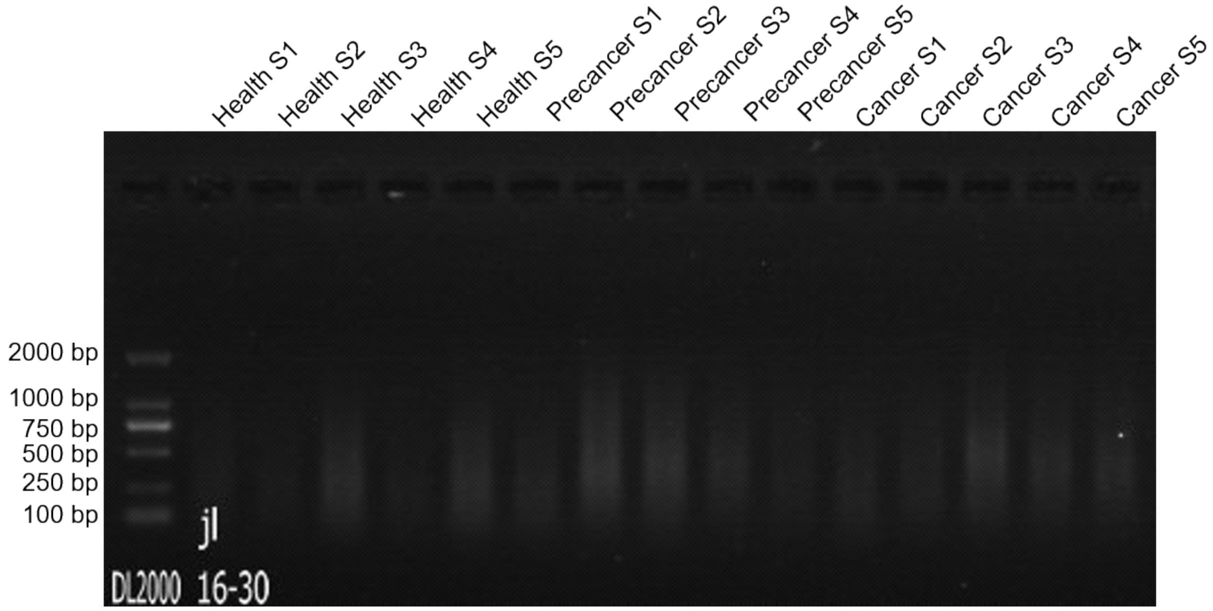


**Figure S6. Agarose gel analysis of the DNA extracted from cervical FFPE tissue samples.** The quality of DNA from 15 cervical related FFPE tissue samples (five from each group) was assessed by analyzing ~100 ng DNA on a 1% agarose gel at 100V for 25 min.
